# Morphoelectric properties of inhibitory neurons shift gradually and regardless of cell type along the depth of the cerebral cortex

**DOI:** 10.64898/2026.03.05.709819

**Authors:** Felipe Yáñez, Fernando Messore, Guanxiao Qi, Nima Dehghani, Hanno S. Meyer, Dirk Feldmeyer, Bert Sakmann, Marcel Oberlaender

## Abstract

How can we understand the enormous diversity of the GABAergic inhibitory neurons in the cerebral cortex? To address this question, we quantify the electrophysiological and morphological properties of inhibitory neurons across the depth of an entire cortical column in the rat barrel cortex. We find properties that shift gradually with the cortical depth of the cell bodies across all inhibitory neurons, regardless of their cell types. By isolating morphoelectric variations from their shifts along the cortical depth, we find that the same simple relationships between morphoelectric properties distinguish between the four main molecular cell types of inhibitory neurons at any cortical depth, and in both the rat barrel cortex and mouse primary visual cortex. We provide converging evidence from dense electron-microscopic reconstructions of inhibitory neurons in the mouse visual cortex, and observe comparable depth-dependent shifts in additional datasets from the mouse primary motor cortex and the middle temporal gyrus of the human cortex. Our findings indicate that two different sources of morphoelectric variations can account for the diversity of cortical inhibitory neurons. The first source is molecular cell type-specific, but cortical depth-independent. The second source is cortical depth-dependent, but affects inhibitory neurons similarly across all cell types. We propose that intrinsic developmental specification vs. extrinsic environmental modulation leads to such a decoupling of inhibitory type-specific properties from gradual shifts of these properties with cortical depth.

## Introduction

The glutamatergic excitatory neurons (EXCs) outnumber by far the GABAergic inhibitory neurons (INHs) in the cerebral cortex (e.g. 80-90% vs. 10-20% in rodents (1)). Although the INHs represent a minority of the cortical neurons, the diversity of their anatomical, functional and neurochemical properties exceeds by far that of the EXCs (2–4), which can be subdivided into rather stereotypical cell types (2, 5, 6). It is indeed believed that the diversity of INHs has arisen through evolution to provide the means by which the cerebral cortex can perform its complex operations (7, 8). In any event, an understanding of the diversity of INHs will be critical to understanding cortical function (7).

Due to their enormous diversity, it remains an ongoing debate how to differentiate cortical INHs into distinct types (9). Thus, there is no consensus on how many types of INHs there are in the cerebral cortex (3, 9, 10). However, anatomically, INHs are distinguished by their somatic, dendritic, and axonal morphologies, including the preferences of their axons to target specific subcellular domains (3, 11, 12). Functionally, INHs are distinguished by their electrophysiological responses to current injections (2, 3, 11). Neurochemically, INHs are distinguished by their expression of different molecules, such as different calcium-binding proteins and neuropeptides (3, 4, 7, 9, 11, 13, 14).

However, INHs with comparable properties in one of these modalities frequently show differences in some of their other properties within the same or the other modalities. Indeed, in each modality, some properties may show continuous variations across INHs, while being otherwise unique in other properties (2, 3, 10). For example, basket cells target specifically the soma and proximal dendrites (15, 16), while chandelier cells innervate the axon initial segment of EXCs (17, 18). Although they are anatomically distinct, these two INH types share some functional properties, since both basket and chandelier cells are classified as fast spiking (FS) neurons with highly variable, yet overlapping firing patterns (19). Molecularly, basket and chandelier cells express both the calcium-binding protein parvalbumin (7). This mix of distinct and continuous variations within and across modalities – morphology, electrophysiology, molecular expression – makes an understanding of the diversity of cortical INHs challenging (3, 4, 7, 9, 10).

The Allen Institute for Brain Science initiated efforts to address this challenge for the primary visual cortex (V1) of mice. This initiative has indeed produced some of the most comprehensive quantifications of cortical INH diversity (2, 10, 20). Analysis of these data from all six cortical layers yielded 13 distinct electrophysiological and 19 distinct morphological clusters (2). Combined clustering of electrophysiological and morphological properties together resulted in 26 morphoelectric (ME-)clusters (2). Analysis of genetic properties, including molecular expression patterns, identified 60 distinct transcriptomic (T-)clusters (2, 20). Studies in other cortical areas also arrived at varying numbers of clusters for electrophysiological, morphological and transcriptomic properties of INHs (4, 21, 22). To resolve these discrepancies between modalities, an integrated analysis of the properties from all modalities was suggested (3, 10) – i.e., such multimodal clusters are defined by maximizing cross-modality congruence. An analysis of mouse V1 INHs using this approach resulted in 28 MET-clusters, which were proposed to constitute a unified definition of INH cell types in the cerebral cortex (10).

An interpretation of such multimodal clusters as cell types remains, however, complex and demanding. In fact, the study in mouse V1 revealed transcriptomic clusters with overlapping morphoelectric properties, as well as electrophysiological and morphological heterogeneity within transcriptomic clusters (10). Similarly, MET-clusters overlapped in electrophysiological properties, but had more distinct morphological properties (10). Consequently, the ability to predict MET-clusters from any single modality is low, and prediction errors vary considerably across clusters and differ for predictions from electrophysiological and morphological properties (10). These limitations demonstrate that an interpretation of multimodal clustering results requires a better understanding of how the properties of INHs relate to one another within and across the modalities.

Here we investigated these relationships by quantifying the electrophysiological and morphological diversity of INHs in the vibrissal-related part of the rat primary somatosensory cortex (vS1), the barrel cortex (23). Facial vibrissae are quintessential sensory organs for rodents (24–26). Tactile information from each vibrissa is processed by an anatomically segregated cluster of neurons in L4 – i.e., the barrel (27). When extrapolating a barrel towards the pia and white matter (WM), a barrel column is considered to provide an anatomical substrate for the major functional unit of the cortex – a cortical column (28, 29). The vibrissal system is therefore widely used as a model for investigating the anatomical and physiological substrates of sensory processing at the cortical level (23). In fact, much of what we know about INHs results from the barrel cortex (30). However, a comprehensive analysis of INH diversity across an entire barrel column has yet to be achieved.

We demonstrate here that INH diversity across the six layers of a barrel column resembles that observed in other cortical areas and species (2, 10, 31–33). In fact, the number of ME-clusters that we identify for rat vS1 – and their distinctive properties – are remarkably similar to those reported for mouse V1 (2). Notably, we show that the clusters can be explained by just two sources of morphoelectric variations. The first source is depth-independent, but type-specific – i.e., at any cortical depth of rat vS1 and mouse V1, the same properties distinguish between the main molecular types of INHs (4, 7). The second source is depth-dependent, but type-unspecific – i.e., such properties shift gradually with cortical depth across INHs, largely irrespective of their electrophysiological, morphological and molecular cell type. Moreover, our analysis of electron-microscopic (EM) reconstructions from mouse V1 (34), as well as of published patch-seq datasets from the mouse primary motor cortex (M1) (32) and the middle temporal gyrus (MTG) of the human cortex (35) reveal comparable depth-dependent shifts in ME-features, while preserving depth-independent relationships that separate INHs by molecular type. In sum, our results indicate that cell type-defining differences between INHs are largely decoupled from gradual shifts of ME-properties along the depth of the cerebral cortex.

## Results

We quantified the electrophysiological and morphological diversity of INHs in the rat barrel cortex **(Fig. 1A)**. For this purpose, we examined the electrophysiological and morphological properties for 13.4% of the INHs that are contained, on average, within a barrel column **(Fig. 1B)** – i.e., for 306 of 2314 INHs (1). We sampled these INHs at any cortical depth by performing whole-cell recordings in brain slices (*in vitro*) from 296 INHs in L2-L6 (36–41), and in intact brains (*in vivo*) from 10 INHs in L1 (42). The distribution of sampled INHs across the cortical depth **(Fig. 1C)** was consistent with the distribution of all INHs in rat vS1 (KS-test: p=0.07) (43). Thus, our sample accounts for the relative occurrence of the INHs at any depth of the barrel column.

**Figure 1:**
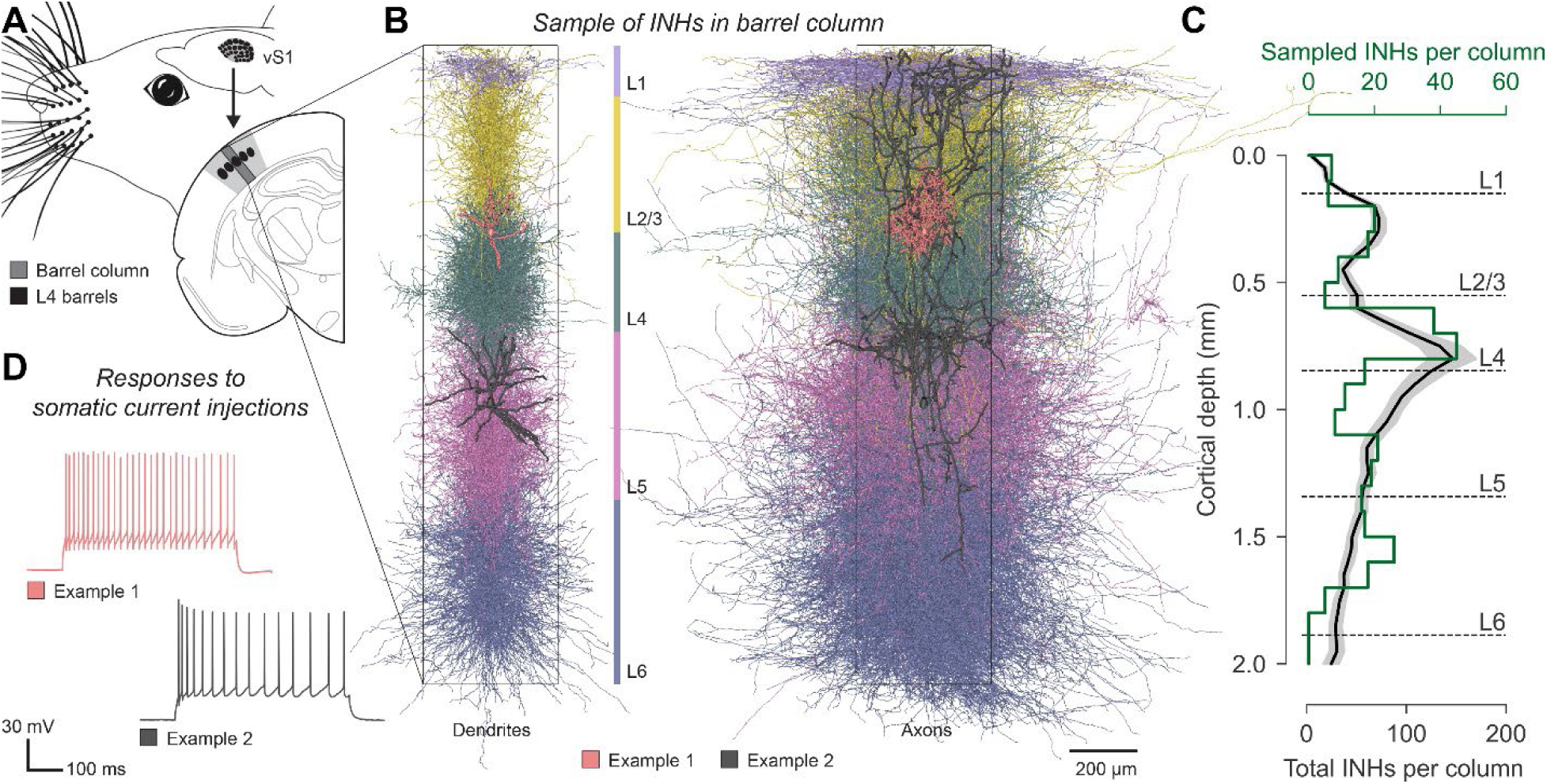
Sampling the morphoelectric diversity of INHs across the depth of a cortical barrel column. **(A)** Schematic of the rat vibrissal system. Tactile information from each facial vibrissa is processed by an anatomically segregated cluster of neurons in layer 4 (L4) – the barrel – of the vibrissal-related part of the primary somatosensory cortex (vS1). By extrapolating a barrel towards the pia and white matter (WM), the barrel column is considered to provide an anatomical substrate for the major functional unit of the cortex – the cortical column. Here we quantified electrophysiological and morphological properties of 306 inhibitory neurons (INHs) that we sampled across the depth of a cortical barrel column. **(B)** We registered each INH to an average barrel column (black box) to identify its soma depth location with 50 µm precision (43). Dendritic (left) and axonal morphologies (right) of INHs are colored by their soma locations in L1-6. The two examples represent a basket cell in L2/3 (red) and a Martinotti cell in L5 (gray). **(C)** We had reported the number and distribution of INHs in entire rat vS1 (1). The hence determined distributions of INHs along the depth of the barrel columns (the black line represents the mean, the gray-shading the 95% confidence interval across the barrel columns of the 24 major vibrissae from 4 rats) were not significantly different from that of the INHs whose morphoelectric properties were sampled here (green; KS-test: p=0.07). **(D)** Action potential (AP) responses to step-currents for the example INHs in panel B. The AP responses of the basket cell (red) and the Martinotti cell (gray) are characteristic for fast-spiking (FS) and non-FS INHs, respectively.

In response to step-current injections, the evoked action potential (AP) patterns varied substantially across our sample of INHs **(Fig. 1D)**. Within each cortical layer, we observed AP patterns that are characteristic for the main electrophysiological INH types (E-types): fast-spiking (FS) versus non-FS cells, and the latter with regular or irregular and adapting or non-adapting AP patterns (3), respectively **(Fig. S1A)**. During the electrophysiological recordings, the INHs were labeled with biocytin for *post hoc* reconstruction of their dendrites and axons **(Fig. 1B)**. These reconstructions showed properties that are characteristic for the main morphological types of cortical INHs (M-types) (3), including basket, chandelier, Martinotti, neurogliaform and bipolar cells **(Fig. S1B)**. Overall, we conclude that our sample comprised the main electrophysiological and morphological types of cortical INHs.

### Sample represents morphoelectric diversity of INHs in a barrel column

Next, we examined to which extent our sample could be considered to represent the full electrophysiological and morphological diversity of INHs across the depth of a barrel column. For this purpose, we modified the multimodal clustering approach (see **Fig. S2** and **Methods** for more details) used by Gouwens et al. to quantify the diversity of INHs in mouse V1 (2, 10).

For each INH, we quantified electrophysiological (n=31, **Table S1**) and morphological features (n=47, **Table S2**) that are considered characteristic of the different INH (sub)types (2, 6, 9, 22, 37, 40, 41, 44–46). Electrophysiological features included parameters such as AP waveforms and inter-spike-intervals (ISIs), as well as changes of these features with stimulus duration (e.g. adaptation of ISIs to longer intervals). Morphological features included parameters such as the distributions of dendrites and axons across cortical layers, the dendritic and axonal pathlengths, as well as the cortical depth of the soma. We z-scored these features **(Fig. 2A)** and discarded ten because of low coefficients of variation (CV < 0.25) or high correlations with other features (Pearson R > 0.95).

**Figure 2:**
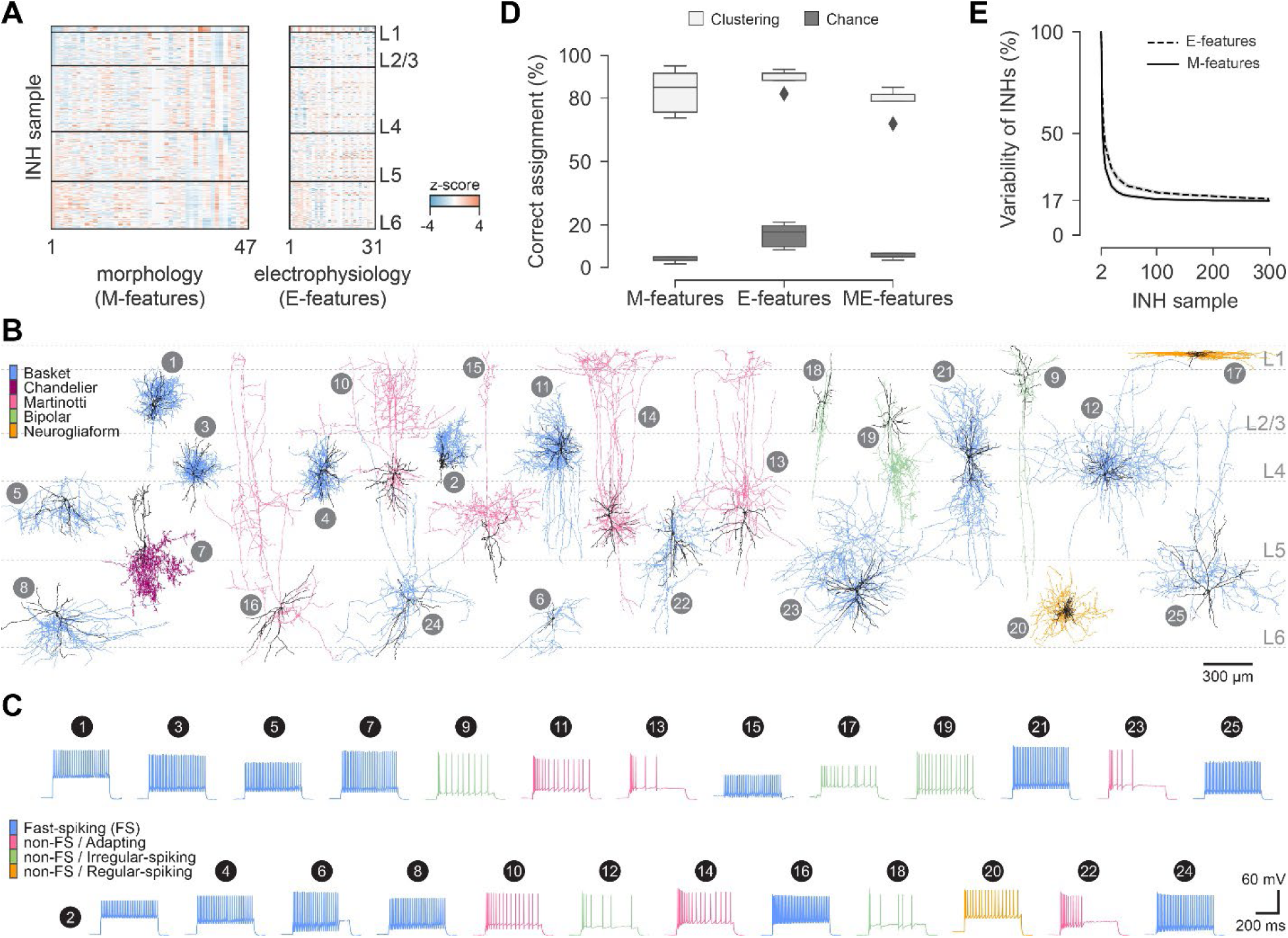
Sample of INHs accounts for the morphoelectric diversity within a barrel column. **(A)** We quantified morphological (M-)features (left, **Table S2**) and electrophysiological (E-)features (right, **Table S1**) for each INH of our sample that were suggested to characterize INHs (2, 6, 9, 22, 37, 40, 41, 44–46). We z-scored these morphoelectric (ME-)features and show them sorted by soma depths of the INHs within the barrel column. **(B)** Example morphologies for each of the 25 ME-clusters. Dendrites are colored in black. Axons are colored by major M-types of INHs. **(C)** Electrophysiology (i.e., AP responses to step-current injections) for the example morphologies from panel B, colored by major E-types of INHs. **(D)** Validation of our final cluster assignment. We trained classifiers on the M-, E- or ME-features of 80% of the INHs and predicted the assignment to clusters for the remaining 20% of the INHs **(Fig. S3)**. Box plots (light grey) represent medians and 25^th^/75^th^ percentiles across five validations and five random shuffles (dark grey). **(E)** Variability of M- and E-features (mean ± 95% CI of principal component variance) as a function of the number of sampled INHs in rat vS1.

Based on the remaining 27 electrophysiological and 41 morphological features, we generated initial cluster assignments (47). We repeated this process with four algorithms: hierarchical clustering with and without connectivity constraints, Gaussian mixture model and spectral clustering. The four algorithms resulted in 60 different instances of how our INH sample could be separated into ME-clusters. Hierarchical clustering with connectivity constraints yielded 12 initial clusters, the other algorithms each yielded 16 initial clusters.

We generated the final cluster assignment by calculating similarity scores between all pairs of INHs based on how often they were grouped together in the 60 initial clusters. We iteratively performed hierarchical clustering, cluster merging and reassignment with the similarity scores. This consensus clustering algorithm terminated by identifying the most robust and hence final cluster assignment for our INHs. Consistent with the study that identified 26 ME-clusters of INHs from mouse V1 (2), our consensus clustering concluded that 25 ME-clusters represent the most robust clustering of INHs from rat vS1 **(Fig. 2B-C)**.

We validated our final cluster assignment and tested the robustness of this validation across seven classifiers **(Fig. S3)**. Independent of the classifier we used, this analysis showed that the percentages for predicting the correct cluster assignment for INHs in our sample from rat vS1 outperforms that reported for the sample of INHs from mouse V1 by a margin of up to 10% **(Fig. 2D, S3)**.

In summary, our study demonstrates that the electrophysiological and morphological diversity in our sample of INHs from rat vS1 is analogous to that observed in other cortical areas and species (2, 10, 31–33). In fact, not only the number of INH clusters, but also their characteristic features, show a remarkable similarity between rat vS1 and mouse V1 (see **Discussion**). Moreover, the robustness of our clusters rivals and even exceeds that reported for mouse V1 (2). These results indicate that our sample accounts for the electrophysiological and morphological diversity of INHs across the depth of a barrel column. In support of this interpretation, we show that the variability of both electrophysiological and morphological features saturates for samples smaller than that analyzed in this study **(Fig. 2E)**.

### Morphoelectric properties of INHs shift gradually with cortical depth

We calculated the Gini impurity index (48), which estimates the relative importance of each feature for cluster assignment **(Fig. S4A)**. This analysis identified the cortical depth of the somata as the main contributor for cluster assignment. To confirm this result, we visualized our sample in a 2D version of the feature space that gave rise to the 25 ME-clusters via a uniform manifold approximation and projection (UMAP) algorithm (49). This visualization confirmed that INHs whose somata are located at the same cortical depth will also be located close to one another in the feature space that accounts for the ME-diversity of INHs in the barrel cortex **(Fig. S4B)**.

Therefore, we examined how variations of other electrophysiological and morphological features are related to cortical depth **(Fig. 3A)**. For this purpose, we unfolded the UMAP along its cortical depth dimension (see **Methods** and **Fig. S4B** for details). As a result, the soma depth gradient from the pial surface to the white matter (WM) was well-aligned with the horizontal axis of the UMAP **(Fig. 3B-C)**. The unfolding enabled us to assess how variations of each ME-feature relate to cortical depth. In essence, by calculating the gradients of features across the unfolded UMAP **(Fig. 3D-G, S4C)**, we revealed whether a feature varies with cortical depth (i.e., horizontal gradient), independent of cortical depth (i.e., vertical gradient), or both (i.e., diagonal gradient). We subsequently refer to the unfolded UMAP as ‘ME-diversity space’ of INHs in the barrel cortex.

**Figure 3:**
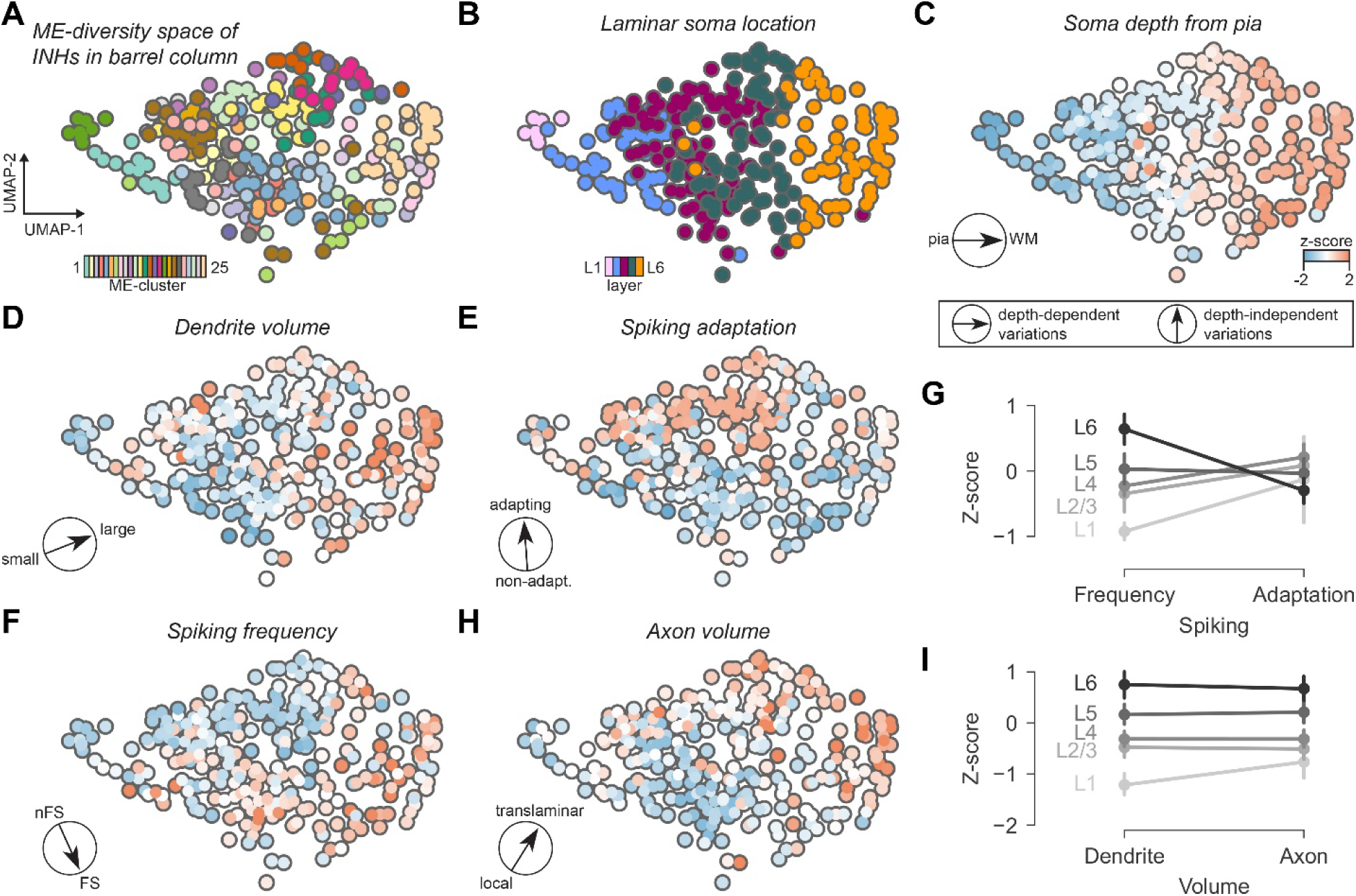
Morphoelectric properties of INHs shift gradually with cortical depth. **(A)** We visualized our sample of INHs in a 2D version of the feature space that gave rise to the 25 ME-clusters via a uniform manifold approximation and projection (UMAP) algorithm and ‘unfolded’ the UMAP along the cortical depth dimension **(Fig. S4B)**, which is the most important feature for cluster assignment **(Fig. S4A)**. We refer to the unfolded UMAP as ME-diversity space of INHs in the barrel column. **(B)** We colored the INHs in the ME-diversity space by their soma location in L1-L6. **(C)** We colored the INHs by their soma depths. The gradient of this feature (black arrow) was aligned with the horizontal axis of the ME-diversity space – i.e., soma depth structures the space from left (pia) to right (white matter, WM). **(D)** We colored the INHs by their dendrite volumes, which showed a nearly perfect horizontal gradient – i.e., this feature varies primarily depending on cortical depth. **(E)** We colored the INHs by their spiking adaptation, which showed a nearly perfect vertical gradient – i.e., this feature varies primarily independent of cortical depth and hence, it structures the space from top (strongest adapting INHs at any depth) to bottom (non-adapting INHs at any depth). **(F)** We colored the INHs by their spiking frequencies, which showed a diagonal gradient – i.e., this feature varies both depending and independent of cortical depth. Like adaptation in panel E, spiking frequency structures the ME-diversity space from top (non-FS cells) to bottom (FS cells). However, like dendritic volumes in panel D, the entire distributions shift gradually to higher spiking frequencies with increasing cortical depth. **(G)** Spiking frequencies shift to higher values from L1 to L6, whereas adaption does not shift (mean ± 95% CI of z-scored E-features). **(H)** We colored the INHs by their axon volumes, which showed also a diagonal gradient – i.e., this feature structures the ME-diversity space also from top (translaminar) to bottom (local). **(I)** Dendrite and axon volumes of INHs shift to higher values from L1 to L6, (mean ± 95% CI of z-scored M-features).

Strikingly, important features for the cluster assignment had nearly perfect horizontal gradients – i.e., they structured the ME-diversity space like cortical depth, and hence from left (pia) to right (WM). For example, INHs show similar variations of dendritic innervation volumes of at any cortical depth, but the distributions shift gradually to larger volumes with increasing cortical depth **(Fig. 3D)**. Other features had nearly perfect vertical gradients – i.e., they structured the ME-diversity space independent of cortical depth, and hence from top to bottom. For example, the adaptation of the spiking frequency during stimulus duration did not increase with cortical depth **(Fig. 3E)**. In contrast, the distributions of this feature were largely the same at any cortical depth: weakly and non-adapting INHs populated the bottom portion of the ME-diversity space, whereas the strongly adapting INHs populated the top portion. Consequently, dendritic innervation volumes increase with cortical depth, regardless of whether spiking patterns of INHs show strong or weak adaptation, or no adaptation at all. Thus, features that characterize INHs can vary independent from one another – i.e., variability across the depth of the barrel cortex (e.g. dendritic volume) is independent from variability within any cortical depth (e.g. spiking frequency adaptation).

For most features we observed diagonal gradients, indicating that such features show variations both within any cortical depth and across cortical depths **(Fig. S4C)**. For example, like adaptation, the spiking frequency distributions of INHs structured the ME-diversity space from top to bottom. At any cortical depth, FS INHs occupied the bottom portion of the ME-diversity space, whereas non-FS INHs occupied the top portion **(Fig. 3F)**. However, in contrast to adaptation, but as for dendritic innervation volumes, the entire distributions of spiking frequencies gradually shifted to higher values with increasing cortical depth **(Fig. 3G)**. In essence, spiking frequencies increase with cortical depth, regardless of whether INHs are FS or not. Another example for a diagonal gradient was the axon innervation volume, which also structured the ME-diversity space from top to bottom, while shifting gradually to higher values with increasing cortical depth **(Fig. 3H)**. At any depth, INHs with axons confined to a single cortical layer (i.e., local cells) occupied the bottom portion of the ME-diversity space, those with axons across all cortical layers (i.e., translaminar cells) occupied the top portion.

Thus, like dendritic volumes and spiking frequencies, also axonal volumes increase with increasing cortical depth, regardless of whether the INHs show a local or translaminar axonal projection pattern **(Fig. 3I)**.

Taken together, variations in the morphoelectric properties of INHs in the barrel cortex can depend solely on cortical depth (e.g. dendrite volume), be fully independent of cortical depth (e.g. adaptation), or depend on both, where the entire distributions that structure the ME-diversity space similarly at any cortical depth shift gradually with increasing cortical depth to higher values (e.g. spiking frequency).

To isolate morphoelectric variations that are independent of cortical depth from the gradual shifts with depth, we grouped INHs based on their variations along the vertical axis of the ME-diversity space (see **Methods** for details). These depth-independent variations in spiking frequency and adaptation, as well as in dendrite and axon volumes, revealed three distinct groups of INHs, which occupied approximately the bottom, center and top thirds of the ME-diversity space **(Fig. 4A)**. Group I (bottom) comprised INHs, which at their cortical depth had the fastest spiking frequencies, most locally confined axons and no adaptation (1-way ANOVA: for all p<10^−6^, **Fig. 4B**). In turn, Group II (top) comprised INHs, which at their cortical depth had the slowest spiking frequencies, most extensive translaminar projecting axons and strongest adaptation (p<10^−11^, **Fig. 4C**). Group III (center third) comprised INHs with spiking frequencies similar to those in Group II, but less elaborate axons and weaker adaptation (p<10^−8^, **Fig. 4D**). Group III also comprised INHs with the smallest dendrites (p<10^−14^, **Fig. 4E**), which we treated as subgroup IIIb, because compared to the INHs with normal sized dendrites (subgroup IIIa), they had less elaborate axons and slower spiking frequencies (p<0.002).

**Figure 4:**
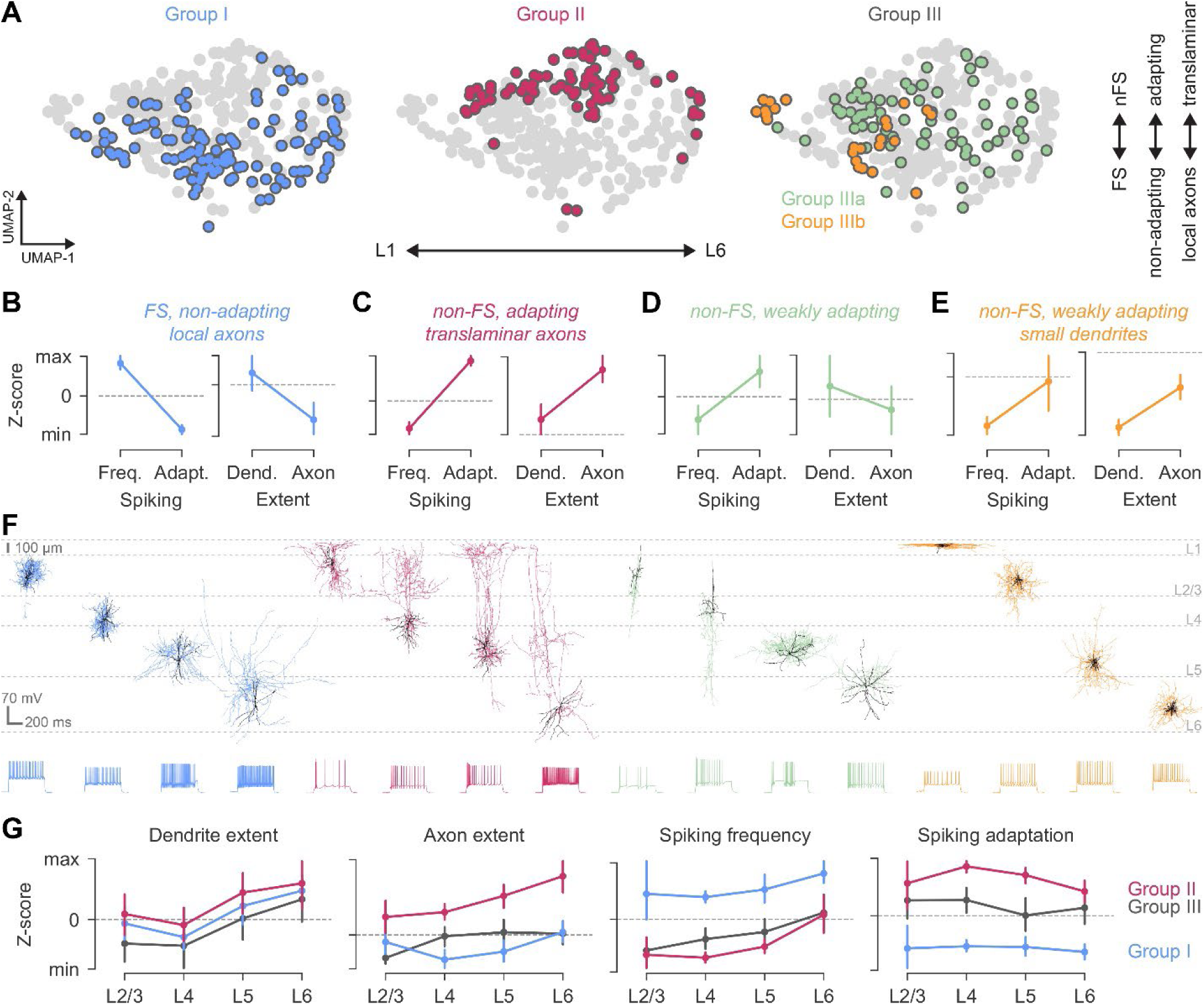
ME-clusters of INHs condense into three depth-independent groups. **(A)** We grouped INHs based on their variations along the vertical axis of the ME-diversity space, to isolate ME-variations that are independent of cortical depth from their gradual shifts with depth (see **Methods** for details). The isolated variability in spiking frequency and adaptation, as well as in dendrite and axon volume, structured the ME-diversity space into three groups, which populated approximately the bottom (blue), center (green and orange) and top thirds of the space (magenta). **(B-E)** Group I (blue) comprised the INHs, which at their respective cortical depths, had the fastest spiking frequencies, most local axons and no adaptation. Group II (magenta) comprised the INHs, which had the slowest spiking frequencies, strongest adaptation and most translaminar axons at their respective cortical depths. Group III comprised INHs with spiking frequencies similar to those in Group II, but weaker adaptation and less elaborate axons. Group III comprised the INHs with the overall smallest dendrites (IIIb, orange), which we treated as a subgroup, because compared to INHs with normal sized dendrites (IIIa, green), they had also less elaborate axons and slower spiking frequencies. **(F)** Morphology and electrophysiology of INH examples for the four major depth-independent groups. Axons are colored by Groups I-IIIa/b, and dendrites are colored in black. **(G)** From left to right: dendritic and axonal extents and spiking frequencies increase with cortical depth similarly in Groups I-III, while these ME-features, as well as adaptation, separate between INHs in Groups I-III at any cortical depth.

Taken together, when gradual shifts of electrophysiological and morphological properties along the depth of the barrel cortex are not considered, the 25 ME-clusters of INHs condense into four depth-independent groups **(Fig. 4F**). Spiking frequencies, as well as dendrite and axon volumes increase similarly with cortical depth in all three groups, while these features, as well as adaptation, separate between the groups at any cortical depth **(Fig. 4G)**. Essentially, Group I separates FS INHs (e.g. basket cells) from non-FS INHs (i.e., Group II/III). In addition, Group II separates strongly-adapting non-FS INHs (e.g. Martinotti cells) from those without or weak adaptation (i.e., Group III). Group IIIa and IIIb separate between non-FS, weakly-adapting INHs with normal sized and very small dendrites (e.g. bipolar versus neurogliaform cells).

### Depth-independent morphoelectric groups reflect the molecular identity of cortical INHs

What is the relevance of these depth-independent groups? Why do some of the main E-types (e.g. FS vs. non-FS) and M-types (e.g. basket vs. bipolar cells) of INHs separate remarkably well at any cortical depth **(Fig. 5A)**? An analysis of the molecular identities of INHs may provide answers to these questions. In fact, ME-features that separate between Groups I-III are reminiscent of those that distinguish between different molecular groups of cortical INHs in rodents. For example, as for Group I, the INHs that express the calcium-binding protein parvalbumin (PV) comprise FS cells, such as basket and chandelier cells (7). As for Group II, the INHs that express the neuropeptide somatostatin (Sst) comprise non-FS strongly adapting cells, such as Martinotti cells (7). As for Group IIIa/b, the INHs that neither express PV nor Sst (double negative: -/-) comprise a heterogeneous set of INHs that splits into (at least) two subgroups (7), those that express the neuropeptide Vip (e.g. bipolar cells) and those that do not (e.g. neurogliaform cells). We therefore hypothesized that ME-variations which distinguish Groups I-IIIa/b at any cortical depth reflect the molecular identities of INHs **(Fig. 5B)**. Subsequently, we provide three lines of evidence in support of this hypothesis.

**Figure 5:**
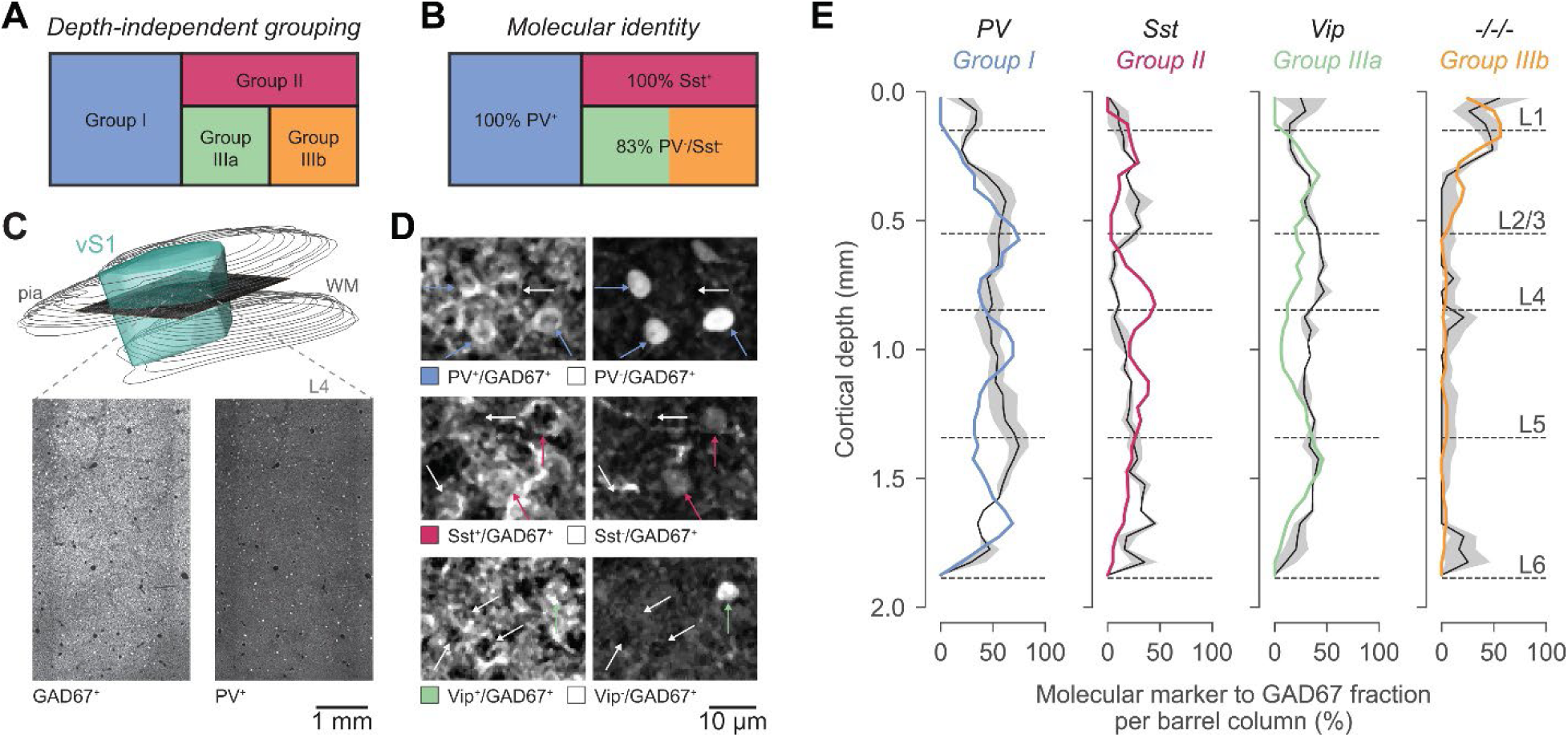
Depth-independent ME-variations of cortical INHs may reflect their molecular type. **(A)** The ME-features that separate between Groups I-IIIa/b are reminiscent of those that distinguish between the major molecular types of INHs (7). Therefore, we hypothesized that ME-variations that separate INHs at any cortical depth into Groups I-IIIa/b may reflect the molecular identities of these INHs – i.e., Group I-III represent the PV^+^, Sst^+^ and PV^−^/Sst^−^ cortical INHs (7), respectively, and Group III is spilt into those that express Vip (IIIa) and those that do not (IIIb). **(B)** For a subset of INHs from our sample (n=14), we used immunohistochemical labeling to identify whether they expressed PV or Sst. The PV^+^ INHs were in Group I (2/2), the Sst^+^ INHs were in Group II (6/6), whereas the double negative INHs were in Group III (5/6). **(C)** We identified all PV^+^, Sst^+^ and Vip^+^ INHs across the entire volume of the rat barrel cortex. For this purpose, we cut 50 μm thick sections tangentially to the barrel cortex from the pial surface to the WM (top), and then double-immunolabeled each section against GAD67 (bottom-left), which is expressed in all INHs (and which also reveals the L4 barrels (1)), and against one of these three molecular markers (bottom-right). **(D)** High-resolution example images show GAD67^+^ INHs (left panels) that co-express PV, Sst or Vip (right panels, colored arrows), or that do not (white arrows). **(E)** Based on GAD67 expression (panel C, bottom-left), we delineated the barrels for the 24 major vibrissae, and hence, we could quantify the distributions of PV^+^, Sst^+^ and Vip^+^ INHs along the depths of these 24 barrel columns **(Table S3)**. The hence measured depth-distributions of PV^+^, Sst^+^, Vip^+^ and triple negative (-/-/-) INHs (black; mean ± SEM, n=6 rats) resemble the respective depth-distributions of Groups I-IIIa/b from our sample of INHs.

First, for a small subset of INHs from our sample (14/306), we used immunohistochemical labeling to identify whether they express PV or Sst (36, 37). PV-positive (PV^+^) INHs were in Group I (2/2), PV-negative (PV^−^) INHs were generally not in Group I (11/12). Sst^+^ INHs were in Group II (6/6), Sst^−^ INHs were not in Group II (0/8). Double negative INHs were generally in Group III (5/6). Thus, immunohistochemical labeling for a small subset of INHs from our sample supports the hypothesis that depth-independent ME-variations reflect the molecular identity of cortical INHs.

Second, we identified all PV^+^, Sst^+^ and Vip^+^ INHs across the rat barrel cortex. For this purpose, we cut 50 μm thick sections tangentially to the barrel cortex from the pial surface to the WM **(Fig. 5C)**, and double-immunolabeled each section against GAD67, which is expressed in all INHs, and against one of these three molecular markers **(Fig. 5D)**. Tangential sectioning enabled us to delineate the barrels for the 24 major vibrissae, and hence, we could quantify the distributions of PV^+^, Sst^+^ and Vip^+^ INHs along the depths of these 24 barrel columns (i.e., by extrapolating L4 barrels to the pia and WM) **(Table S3)**.

We reported previously that, on average, a barrel column comprises 2314 ± 279 INHs (i.e., GAD67^+^ cells; N=4 rats (1)). In this study, we observed 2332 ± 222 INHs (N=3 rats) with a distribution across the depth of a barrel column that was consistent to that from our previous report (two-tailed t-test: p=0.97). Also, the number of PV^+^ and Sst^+^ INHs per cortical layer matched with previous estimates (3, 36, 37, 40, 41, 50). We demonstrate here that 40% of the INHs in a barrel column are PV^+^ (928 ± 119; average across 24 columns, N=2 rats). The other two molecular markers account each for ∼20% of the INHs (Sst^+^: 430 ± 64; Vip^+^: 544 ± 74, N=2 rats). Thus, ∼20% of the INHs in a barrel column were triple negative (-/-/-: 430 ± 92, N=6 rats), which supports previous estimates that Vip^+^ and Vip^−^ (i.e., -/-/-) INHs form equally large subgroups (4, 7).

The distributions of the respective molecular groups resembled closely those of the depth-independent ME-groups **(Fig. 5E)**. Indeed, Group I accounted well for the distribution of PV^+^ INHs (cosine similarity: S=0.95). Group II accounted well for the distribution of Sst^+^ INHs (S=0.81). Group IIIa and IIIb accounted well for the distributions of Vip^+^ and -/-/- INHs (S=0.92 and S=0.99), respectively. Thus, immunohistochemical labeling across the barrel cortex supports the hypothesis that depth-independent ME-variations reflect the molecular identity of cortical INHs.

Third, we repeated our analysis for INHs from mouse V1. Analogous to how we had collected our sample of INHs from the rat barrel cortex, Gouwens et al., had performed whole-cell recordings and biocytin filling in brain slices from INHs in L1-L6 of mouse V1 (2, 10). Following the recordings, the nucleus was removed to analyze transcriptomic profiles. This patch-seq approach revealed the molecular identity for 466 recorded and reconstructed INHs as PV^+^, Sst^+^, Vip^+^ or Lamp5^+^ (i.e., lysosomal-associated membrane protein 5), the latter being a marker for a subset of Vip^−^ INHs, such as neurogliaform cells in cortical layer 1 (10, 20).

We extracted the ME-features that had separated the INHs in the rat barrel cortex (i.e., vS1) into the depth-independent Groups I-IIIa/b **(Fig. 6A)** for the 466 mouse V1 INHs **(Fig. 6B)**. PV^+^, Sst^+^ and Vip^+^ INHs showed similarly broad distributions of dendrite extents, and at any cortical depth, while the distributions shifted with cortical depth to larger values. Lamp5^+^ INHs had generally the smallest dendrites. Similarly, the distributions of axon extents shifted with cortical depth to larger values, while separating INHs at any cortical depth into local (mostly PV^+^) and translaminar cells (mostly Sst^+^ or Vip^+^). EM reconstructions of mouse V1 (34) support these results by demonstrating that the gradual shifts with cortical depth of dendritic and axonal extents are observable, regardless of cell type **(Fig. 6C)**, within the same mouse **(Fig. 6D)**. Moreover, spiking frequency and adaptation separated also the INHs in mouse V1 at any cortical depth into FS, non-adapting (generally PV^+^) and non-FS, strongly/weakly-adapting (generally Sst^+^ / Vip^+^ or Lamp5^+^) cells. Spiking frequency distributions shifted with cortical depth to higher frequencies, whereas adaptation distributions did not. Thus, variations across and within cortical depths of the ME-features that distinguish INHs in rat vS1 into depth-independent Groups I-IIIa/b were remarkably similar for INHs in mouse V1.

**Figure 6:**
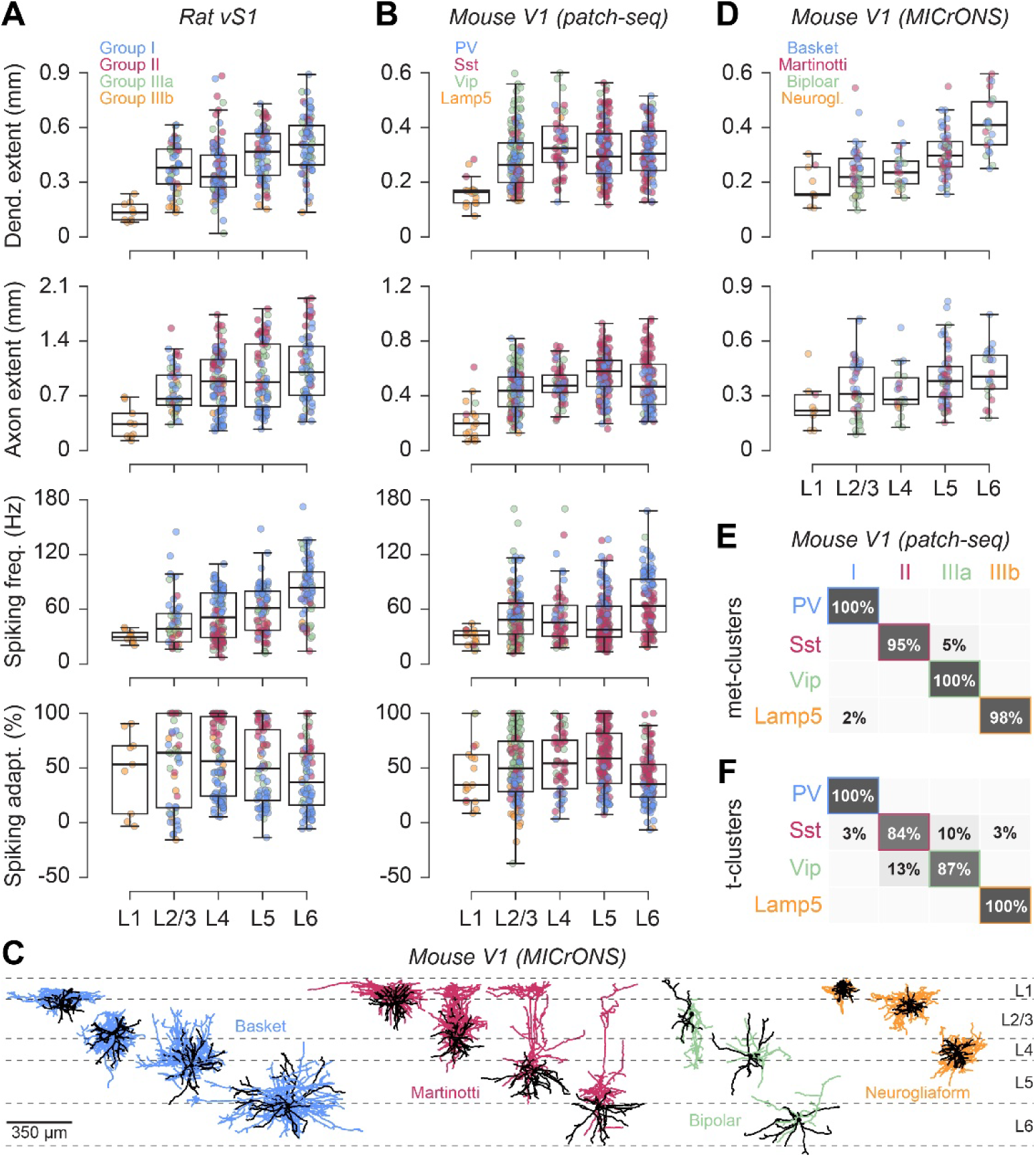
Depth-independent ME-variations of cortical INHs predict their molecular type. **(A)** ME-variations within each cortical layer of the features that had separated the INHs in the rat vS1 into the depth-independent Groups I-IIIa/b. Distributions of dendritic and axonal extents, and those of spiking frequencies, shift with cortical layer to larger values, whereas adaptation distributions do not. **(B)** Same as in panel A for a dataset that was used to classify the electrophysiology and morphology, as well as transcriptomic profiles of 466 INHs across the depth of the mouse primary visual cortex (V1) (2, 10). The variations of these ME-features within and across cortical layers of mouse V1 were remarkably similar to those in rat vS1. Colors denote the molecular type of each INH as derived from their transcriptomic profiles, and match the scheme for the corresponding depth-independent Groups I-IIIa/b in panel A. **(C)** Examples of INH dendrites (black) and axons (colored by morphological cell type and matching the scheme for the corresponding depth-independent Groups I-IIIa/b in panel A) from a dense electron-microscopic (EM) reconstruction of mouse V1 (i.e., MICrONS dataset (34)). **(D)** Distributions of dendritic and axonal extents for the different INH cell types from panel C show that these morphological features shift gradually along the depth of mouse V1 within the same animal **(E)** Based on the criteria identified for rat vS1, we assigned the INHs from mouse V1 to Groups I-IIIa/b. These depth-independent ME-variations predicted the molecular identities for 453 of the 466 INHs from mouse V1. **(F)** Even when INHs are clustered solely by their gene expression profiles – i.e., transcriptomic clusters (20), the depth-independent variations of the four ME-features in panel B still predicted the molecular types for 89% of the INHs in mouse V1..

Therefore, we assigned INHs from mouse V1 to Groups I-IIIa/b based on the relationships we identified in rat vS1. These depth-independent ME-groups predicted the molecular identity for 453 of the 466 INHs from mouse V1 **(Fig. 6E)**. All PV^+^ INHs were in Group I (88/88), 95% of the Sst^+^ INHs in Group II (219/231), all Vip^+^ INHs in Group IIIa (103/103), and 98% of the Lamp5^+^ INHs in Group IIIb (42/43). When we considered variations between INHs in their gene expression profiles alone – i.e., transcriptomic clusters (20), the depth-independent ME-groups still predict molecular identities for 89% of the INHs in mouse V1 **(Fig. 6F)**. These results provide direct support for the hypothesis that depth-independent ME-variations reflect the molecular identity of INHs. In any event, we demonstrate for mouse V1 that at any cortical depth, a higher spiking frequency in the respective distribution separates the PV^+^ from the Sst^+^, Vip^+^ and Lamp5^+^ INHs. In addition, a stronger adaptation separates at any cortical depth the Sst^+^ from the Vip^+^ and Lamp5^+^ INHs, and a very small dendrite extent separates the Lamp5^+^ from the Vip^+^ INHs.

### Morphoelectric properties of INHs shift gradually with cortical depth across areas and species

Dendritic and axonal extents, as well as spiking frequencies, showed remarkably similar gradual shifts along the depth of both rat vS1 and mouse V1. When these shifts were decoupled from the remaining variability of these ME-features, the same relationships between dendritic and axonal extents, spiking frequencies and adaptation – the latter did not shift with cortical depth – predicted whether INHs at any cortical depth express PV, Sst, Vip or Lamp5. Our analysis of two additional patch-seq datasets, one from mouse M1 (32) and one from human MTG (35), suggest that these simple depth-independent relationships between ME-features and molecular type generalize beyond the primary sensory areas of the rodent cerebral cortex.

Although the datasets from mouse M1 and human MTG showed sampling biases for the superficial layers **(Fig. 7A)**, we observed significant shifts with cortical depth of dendritic **(Fig. 7B)** and axonal extents **(Fig. 7C)**, and in spiking frequencies **(Fig. 7D)** that were comparable to those in rat vS1 and mouse V1. Moreover, in all four datasets, the distributions of these three ME-features across and within any cortical depth showed similar relationships with the INH’s molecular type. PV^+^, Sst^+^ and Vip^+^ INHs had similarly broad distributions of dendritic extents, and at any cortical depth, but the distributions shifted with cortical depth to larger values. Lamp5^+^ INHs had generally the smallest dendrites. Axonal extent distributions shifted with cortical depth to larger values, while separating INHs at any cortical depth into local (mostly PV^+^) and translaminar (mostly Sst^+^ or Vip^+^) cells. Spiking frequency distributions shifted with cortical depth to higher frequencies, while separating INHs at any cortical depth into FS (mostly PV^+^) and non-FS (mostly Sst^+^ / Vip^+^ or Lamp5^+^) cells. The slopes of these shifts with cortical depth differed, however, between ME-features within the same cortex area, and for the same ME-feature between cortex areas **(Fig. 7E)**. Finally, when the shifts along the cortical depth are neglected, the relationships between these three ME-features separate INHs in rat vS1, mouse V1, mouse M1 and human MTG equally well according to their molecular type **(Fig. 7F)**.

**Figure 7:**
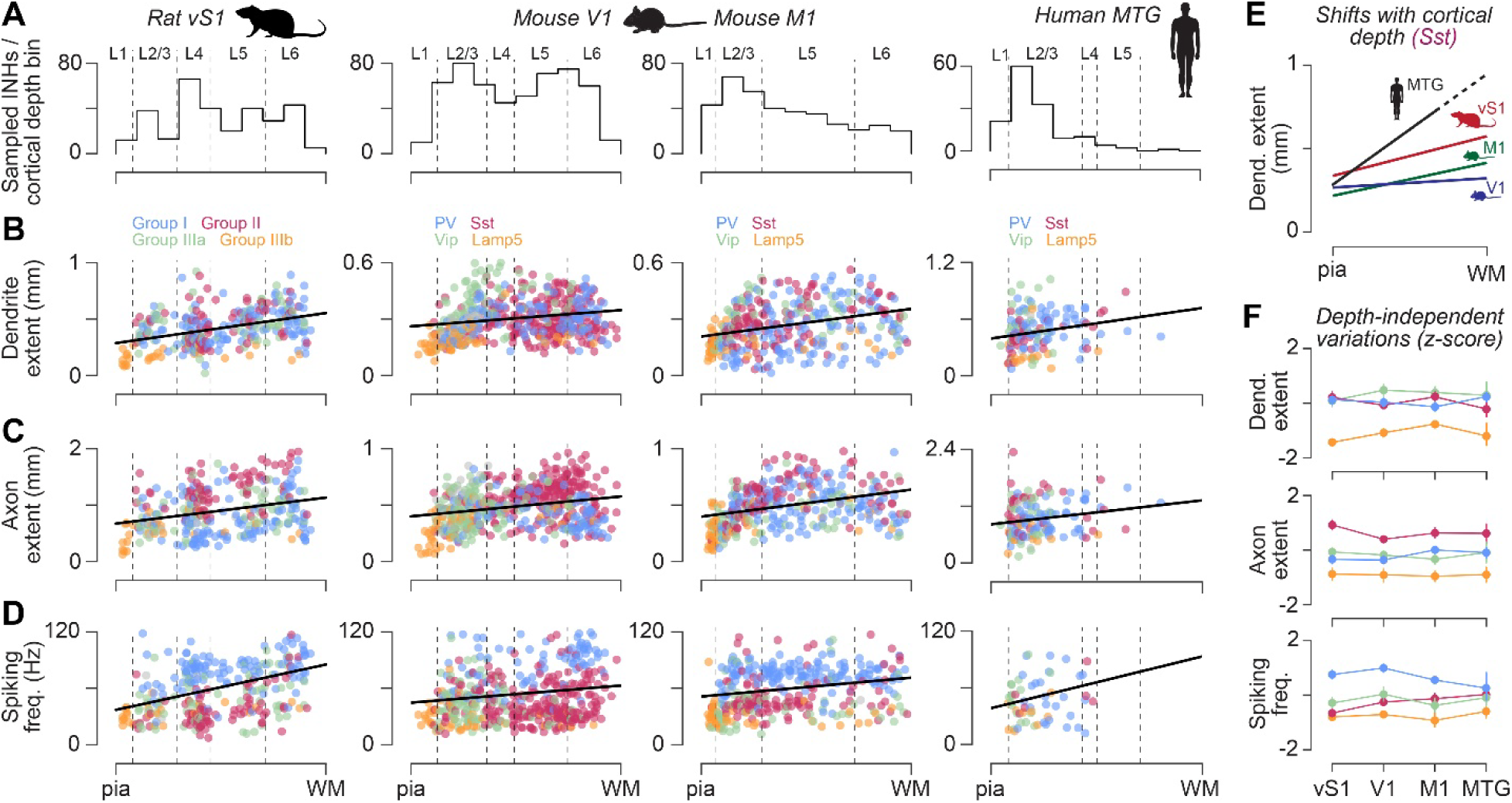
ME-properties of INHs shift similarly with cortical depth across areas and species. **(A)** We compared our sample of INHs from rats vS1 with those reported for mouse V1 (10), mouse primary motor cortex (M1 (32)) and the middle temporal gyrus (MTG) of the human cortex (35). **Note**: in contrast to the present study of INHs sampled from rat vS1, INH samples from mouse V1, M1 and human MTG do not account for the occurrences of INHs at each cortical depth. It is therefore possible that the distributions of ME-features in panels B-E are affected by different sampling biases. **(B)** Dendritic extents shift significantly with cortical depth across INHs in rat vS1 (n=302, p<10^−11^), mouse V1 (n=528, p<10^−5^), mouse M1 (n=370, p<10^−8^) and human MTG (n=140, p<0.05). **(C)** Axonal extents shift significantly with cortical depth across INHs in rat vS1 (n=306, p<10^−6^), mouse V1 (n=528, p<10^−8^), mouse M1 (n=370, p<10^−9^) and human MTG (n=140, p<0.05). **(D)** Spiking frequencies shift significantly with cortical depth across INHs in rat vS1 (n=303, p<10^−9^), mouse V1 (n=466, p<0.005) and mouse M1 (n=363, p<0.001). **Note**: the small sample size of INHs from human MTG did not allow us to draw firm conclusions (n=59, p=0.14). **(E)** The slopes of the gradual shifts with cortical depth generally differed between ME-features within the same cortex area (panels B-D), as well as for the same ME-feature between cortex areas, as exemplified here for the dendritic extents of Sst^+^ INHs in rat vS1 (n=76, slope=0.24, p<0.01), mouse V1 (n=251, slope=0.06, p=0.06) and M1 (n=108, slope=0.24, p<10^−4^) and human MTG (n=33, slope=0.66, p<10^−3^). **Note:** such differences may reflect the different cortical thickness and different laminar organization of these cortical areas, and hence differences in the local environments of the INHs in each cortical layer (see Fig. 8). **(F)** The variations of dendritic and axonal extents, as well as of spiking frequencies, that are independent of cortical depth (i.e., here z-scored across all cortical depths) separate equally well between molecular types of INHs in rat vS1, mouse V1 and M1, and human MTG.

## Discussion

It is well known that INHs in the cerebral cortex display an enormous diversity in their electrophysiological and morphological properties (2–5, 9, 10, 30, 33, 44). It has been demonstrated that some of these ME-variations align with the expression of distinct molecular markers (3, 7), and hence with the transcriptomic profiles of INHs (10, 20). Furthermore, some of these ME-variations were found to correlate with soma location in a specific cortical area and layer therein (2–4, 9, 10, 30). Consequently, when INHs are clustered by their ME-properties, the resulting clusters typically comprise INHs that share molecular identity and layer location (10, 22). Such clusters were suggested to represent a unified definition for INH types, thereby connecting the electrophysiological and morphological properties of INHs with their transcriptomic profiles (10). However, our results indicate two major alterations to this perspective. First, we found electrophysiological and morphological properties that shift gradually with depth across INHs, largely regardless of their morphological, electrophysiological and molecular types. Second, we demonstrate that, at any cortical depth of rat vS1 and mouse V1 – and likely of mouse M1 and human MTG – the same simple relationships between ME-properties distinguish between the four main molecular types of INHs. These findings suggest that depth-independent ME-variations that distinguish between molecular types are decoupled from depth-dependent, and hence layer-specific shifts of these properties – e.g. the spiking frequency increases with cortical depth across INHs, but at any cortical depth, INHs with the highest spiking frequencies in the distribution represent PV^+^ neurons, while the slowest spiking INHs in the distribution represent Sst^+^ neurons.

Below, we will discuss implications of decoupling molecular type- from cortical depth-specific properties of INHs. However, before proceeding further, we will review some of the technical features of our approach. Our findings are based on an analysis of a comprehensive dataset of INHs that we sampled across the depth of a barrel column in rat vS1. We demonstrate that our sample accounts for both the variation of the numbers and the morphoelectric diversity of these INHs across the depth of a barrel column. Consequently, our sample is not biased towards INHs from a particular cortical layer, or from a specific electrophysiological or morphological type.

Cluster analysis supports the conclusion that our sample represents the ME-diversity of cortical INHs across the depth of a barrel column. In fact, the 25 ME-clusters that we identified from our sample of INHs in rat vS1 resembled closely the 26 ME-clusters that were reported from an even larger sample of INHs in mouse V1 (2). Notably, the robustness of our cluster assignment in vS1 exceeded that in V1 (2, 10). Overall, our dataset provides a quantitative account for the electrophysiological and morphological diversity of INHs in rat vS1, and thereby complements similarly comprehensive datasets from other cortical areas and species (2, 10, 31–33).

One clear demonstration of the analysis of our clustering results is that cortical soma depth contributes the most to the ME-diversity of INHs. Essentially, INHs whose somata occupy the same cortical depth are also close to one another in the ME-diversity space – and hence cluster together. This finding is consistent with previous reports, irrespective of whether these studies clustered morphological or electrophysiological properties, or both (2, 10, 22). Therefore, it is believed that depth- and hence layer-specific properties could be used as a distinguishing criteria for defining cortical INHs as different neuronal cell types (10).

However, we found that properties, which gave rise to the ME-clusters, change with depth across all cortical INHs, regardless of their cell types. For example, the spiking frequency shows similarly broad distributions across INHs at any cortical depth of both rat vS1 and mouse V1. This depth-independent variability clearly separates the FS from the non-FS INHs. However, the entire spiking frequency distributions shift gradually towards higher frequencies with increasing cortical depth **(Fig. 3F-G, 4G, 6A-B)**. This depth-dependent, but type-independent variability clearly distinguishes FS from non-FS INHs into depth-specific clusters – i.e., each cortical layer may ultimately comprise a cluster of FS and non-FS INHs, but these splits into layer-specific clusters results from electrophysiological variations that are neither specific to FS nor to non-FS INHs. Incorporating additional ME-properties that shift gradually with cortical depth (e.g. dendritic or axonal volumes) will consequently split both the FS and non-FS INHs into further layer-specific subclusters. This insight is particularly relevant for classification approaches that seek to incorporate an ever expanding set of multimodal data, e.g. by clustering morphoelectric and transcriptomic properties jointly with synaptic connectivity patterns (51). In essence, when interpreting clustering results, it is critical to consider whether the underlying properties show depth-dependent variations that occur similarly across all cortical INHs.

Another clear demonstration of our analysis emerged when we isolated variations of ME-properties from their depth-dependent gradual shifts. The 25 ME-clusters identified in rat vS1, as well as the 26 ME-clusters identified in mouse V1, were condensed into four depth-independent Groups I-IIIa/b. Strikingly, these four ME-groups correspond to the main molecular types of cortical INHs – i.e., PV, Sst, Vip and Lamp5. At any cortical depth, Group I separates the FS from the non-FS INHs (Group II and IIIa/b), and represents PV^+^ INHs. Group II separates the strongly adapting non-FS INHs from the weakly/non-adapting non-FS INHs (Group IIIa/b), and represents Sst^+^ INHs. Groups IIIa and IIIb differentiate between the weakly/non-adapting non-FS INHs with normal-sized and small dendrites, and respectively represent PV^−^/Sst^−^ INHs that express Vip or do not (e.g. Lamp5).

### Intrinsic versus extrinsic influences on morphoelectrical variations of cortical INHs

Overall, we find that the clusters of cortical INHs in both rat vS1 and mouse V1 are the result of two different sources of morphoelectric variations. The first source of variations is depth-independent, but type-specific – i.e., at any cortical depth, the same variations separate between PV^+^, Sst^+^ and PV^−^/Sst^−^ INHs, as well as between Vip^+^ and Vip^−^ INHs within the double-negative group. The second source of variations is depth-dependent, but type-unspecific – i.e., ME-properties shift with cortical depth across INHs, regardless of their electrophysiological, morphological and molecular types – e.g. spiking frequencies, as well as dendritic and axonal innervation volumes increase with cortical depth.

It is indeed remarkable that the same simple relationships between ME-properties can predict the molecular identities of INHs at any cortical depth, and in both rat vS1 and mouse V1. Another way to predict the molecular identities of INHs, independent of their position within a specific cortical area and cortical layer therein, is their place and time of origin during embryological development (52, 53). The majority of cortical INHs originates from progenitor zones of the ganglionic eminences in the ventral telencephalon, from which they migrate to cortex and subsequently integrate horizontally into the different cortical layers. Progenitors from dorsal and ventral regions within the medial ganglionic eminence (MGE) generate the Sst^+^ and PV^+^ cortical INHs, respectively, during overlapping but distinct temporal windows (54, 55). In contrast, the PV^−^/Sst^−^ cortical INHs, including both Vip^+^ cells and neurogliaform cells, are derived from the caudal ganglionic eminence (CGE) or the preoptic region later during development (56–59). The different spatial and temporal patterns of neurogenesis establish transcriptional programs that specify broad INH identity categories and have been found to predict several key morphoelectric features of cortical INHs (4, 13).

Considering these features of cortical INH development together with our results, we propose that the depth-independent morphoelectric variations of cortical INHs **(Fig. 8A)** – which we found here separate the PV^+^, Sst^+^ and PV^−^/Sst^−^ INHs at any cortical depth of both rat vS1 and mouse V1 – reflect, at least in part, intrinsic programs established through cell type specification at their distinct embryological origins (52, 60). Thus, despite the substantial heterogeneity of the INHs within each group – e.g. Group I comprises both basket and chandelier cells, intrinsic developmental programs may determine the broad categories of INH terminal phenotypes as FS versus non-FS, non-adapting versus weakly versus strongly adapting, and local versus translaminar projecting INHs **(Fig. 8B)**.

**Figure 8:**
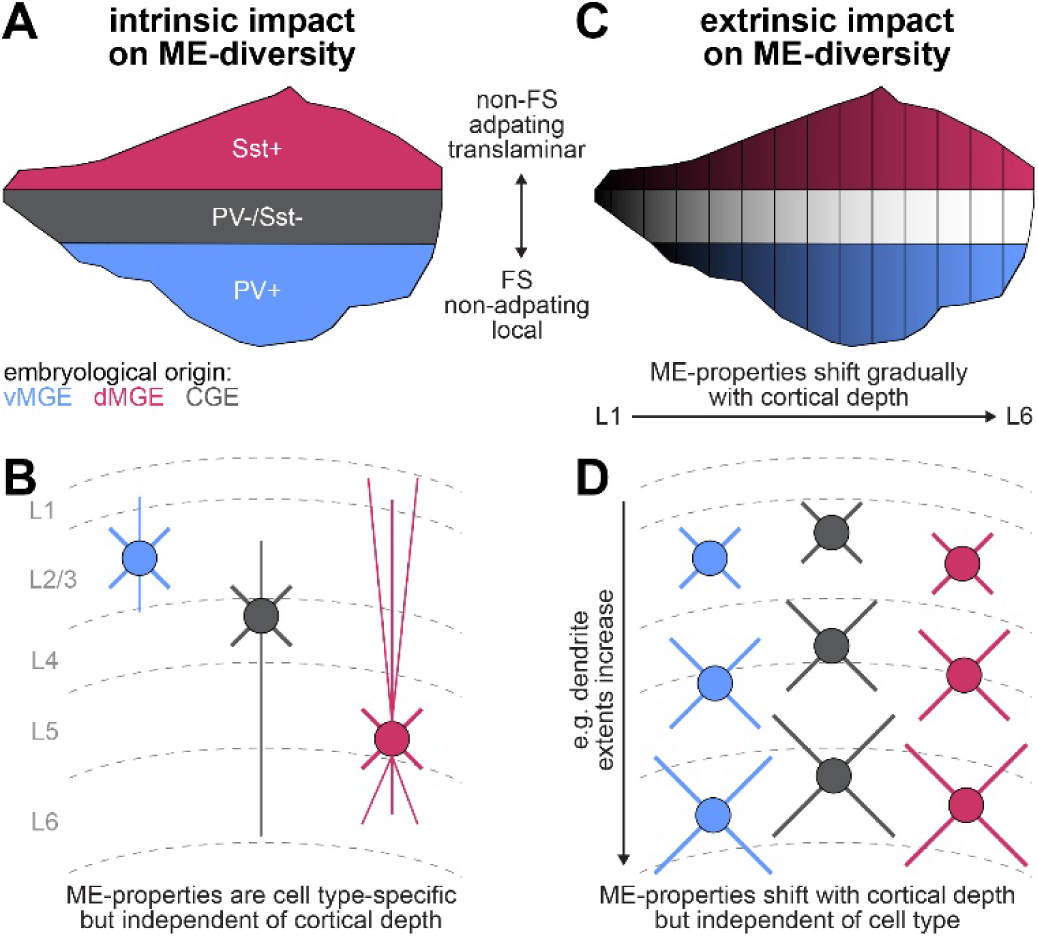
Proposed model to account for the ME-diversity of cortical INHs. The ME-diversity space of cortical INHs in both rat vS1 and mouse V1 is structured by two sources of ME-variations. **(A)** The first source is depth-independent, but type-specific – i.e., at any cortical depth, the same ME-variations separate between the PV^+^, Sst^+^ and PV^−^/Sst^−^ INHs. These molecular identities reflect the origins of cortical INHs during embryological development (52, 53). PV^+^ and Sst^+^ INHs originate from ventral (v) and dorsal (d) regions of the medial ganglionic eminence (MGE), respectively. The PV^−^/Sst^−^ INHs are derived from the caudal ganglionic eminence (CGE), including Vip^+^ and Vip^−^ INHs (56–59). **(B)** We propose that the depth-independent ME-variations of cortical INHs reflect these intrinsic programs established through cell type specification at their distinct embryological origins (52, 60). Thus, intrinsic programs may determine the broad terminal phenotypes as FS vs non-FS, non-adapting vs weakly vs strongly adapting, and local vs translaminar projecting cortical INHs. **(C)** The second source is depth-dependent, but type-unspecific – i.e., ME-properties shift with depth across INHs, regardless of their molecular types. These shifts may hence not be determined by intrinsic programs, but instead by extrinsic factors encountered in the different local environments of each layer during migration and circuit integration (61–63). **(D)** We propose that the gradual shifts of ME-properties with cortical depth reflect, at least in part, such extrinsic factors (45). Thus, extrinsic factors may modulate the terminal phenotypes of cortical INHs by shifting spiking frequency, as well as dendrite and axon volumes to typically higher values with increasing cortical depth.

However, our finding that properties of cortical INHs shift gradually with cortical depth, regardless of their molecular types, suggests that these depth-dependent ME-variations may not be exclusively determined by intrinsic programs. Indeed, extrinsic factors encountered during migration and circuit integration also contribute substantially to both the terminal electrophysiological and morphological properties of cortical INHs (61–63). For example, neuronal activity and experience modulate the electrophysiological features of PV^+^ basket cells by adjusting expression levels of Kv2.1 potassium channels (62). Similarly, neuronal activity and molecular mechanisms that control neurite growth and branching may modulate morphological features of PV^+^ chandelier cells by adjusting axonal innervation volumes to their cortical location (63).

Considering these features of cortical INH development together with our results, we propose that gradual shifts of morphoelectric properties **(Fig. 8C)**, which we found here occur similarly across all INHs, reflect, at least in part, extrinsic factors present in the different local environments of each cortical layer (45). A recent study demonstrated indeed the presence of distinct, pyramidal neuron-driven mechanisms that regulate the diversity and integration of INHs into the local circuitry, and conditional on molecular type (64). Thus, extrinsic factors may modulate the terminal phenotypes of all types of INHs by gradually shifting spiking frequency, as well as dendritic and axonal volume distributions with cortical depth **(Fig. 8D)**. However, for any particular electrophysiological and morphological property, the respective contribution of intrinsic developmental specification versus extrinsic environmental modulation remains to be determined experimentally, though both factors are likely to operate hierarchically to generate the observed diversity of cortical INHs (57, 60).

Taken together, our findings suggest that two complementary sources may account for the enormous ME-diversity of cortical INHs. First, depth-independent ME-variations separate between the main molecular INH types, and may hence reflect the impact of intrinsic factors on broad terminal phenotypes. Second, depth-dependent ME-variations affect all INHs similarly, and may hence reflect the impact of extrinsic, probably intracortical, factors on layer-specific terminal phenotypes. Such decoupling of molecular type- from depth-specific variations may generalize to other properties that were not investigated here. In fact, the impact of extrinsic factors is not limited to the ME-properties, but extends to the synaptic organization and expression of neurochemical markers of cortical INHs (63). Thus, decoupling depth-independent variations from depth-dependent shifts may provide a general strategy to identify which properties of cortical INHs are regulated by intrinsic developmental specification versus extrinsic environmental modulation.

## Materials and Methods

### Animal experiments

All experiments were performed in accordance with the animal welfare guidelines of the Max Planck Society, and with approval by the local German authorities and the Institutional Animal Care and Use Committee. For quantifying the ME-diversity of INHs in L1 of vS1 (42), male and female Wistar rats aged 25-35 days (m/f, P25-P35) were anesthetized with urethane (i.p.; 1.6–2 g/kg body weight). A 2-3 mm-wide craniotomy was opened above vS1 and functional maps of principal and surround whiskers were determined by intrinsic optical imaging (IOI). Neurons located in L1 of the barrel column corresponding to the principal whisker were targeted for whole-cell recordings using 2p microscopy (excitation wavelength, 872 nm; laser model Mai Tai HP; Spectra-Physics). Pipettes were filled with (in mM) K-gluconate 135, 4-(2-hydroxyethyl)-1-piperazineethanesulfonate (Hepes) 10, phosphocreatine-Na 10, KCl 4, ATP-Mg 4, GTP-Mg 0.3, and 0.3–0.5% biocytin, and pipettes had and open resistance of 5-7 MΩ. Fluorescent dye Alexa Fluor 594 (25–50 µM) or OGB-1 (100 µM) was added to visualize the pipette and the patched neuron. Membrane potentials during somatic step current injections (see (42) for details) were recorded using Axoclamp 2-B or MultiClamp 700B amplifiers (Axon Instruments), and digitized using a CED power1401 data acquisition board (Cambridge Electronic Design). Rats were transcardially perfused with 0.1M phosphate buffer (PB) followed by 4% paraformaldehyde (PFA). Brains were removed and post-fixed with 4% PFA for 8 to 12 hours, transferred to 0.1 M PB and stored at 4°C. The cortex was cut tangentially to vS1 (45°) in 100 μm thick consecutive sections (i.e., from the pia to the white matter) and stained for cytochrome oxidase and biocytin (Vectastain ABC-kit). For quantifying the ME-diversity of INHs in L2-L6 of vS1, Wistar rats (m/f, P17-P35) were deeply anaesthetized with isoflurane and decapitated. The brain was removed and transferred to a preparation solution (∼4 °C, 4 mM MgCl_2_, 1 mM CaCl_2_), bubbled continuously with carbogen gas (95/5% O_2_/CO_2_). About 300 or 350 μm thick thalamocortical or semi-coronal slices of vS1 were cut with a vibratome (DTK-1000, Dosaka, Japan or Slicer HR-2; Sigmann Elektronik, Germany). The slices were incubated for ∼1 h (at 21-24 °C) in the solution and subsequently placed in the recording chamber. Neurons were visualized under an upright microscope with a 40× water immersion/0.80 NA objective (Olympus, Germany) under infrared differential inference contrast microscopy. During whole-cell recordings, slices were continuously perfused with artificial cerebrospinal fluid containing (in mM): 125 NaCl, 25 D-glucose, 25 NaHCO3, 2.5 KCl, 2 CaCl2, 1.25 NaH2PO4 and 1 MgCl2, bubbled with carbogen gas, and maintained at a temperature of 31-33 °C. Patch pipettes were filled with a solution containing (in mM): 135 K-gluconate, 4 KCl, 10 HEPES, 10 phosphocreatine, 4 Mg-ATP, and 0.3 GTP (pH 7.4, 290–300 mOsm) with 5 mg/ml biocytin (13.4 mM). Membrane potentials during somatic step current injections were recorded by an EPC10 amplifier (HEKA, Germany) by using patch pipettes of 5-7 MΩ resistance on neurons located at least 50 μm below the slice surface (see (36–41) for details). After the recordings, slices were fixed in 4% PFA (12.9 mM) in 100 mM PB for at least 24 h at 4 °C, treated with 3% H_2_O_2_ (29.4 mM) solution in PB for about 20 min, rinsed repeatedly using 100 mM PB, and finally stained at room temperature for cytochrome oxidase and biocytin (Vectastain ABC-kit). For quantifying distributions of PV^+^, Sst^+^ and Vip^+^ INHs across entire vS1, Wistar rats (m, P28-35) were deeply anesthetized with urethane, transcardially perfused and brains were removed as described above. The cortex was cut tangentially to vS1 into 50 μm thick consecutive sections from the pial surface to the white matter (40-48 sections). The sections were immunolabeled twice: first with antibodies against PV, Sst, or Vip, and then against GAD67. In the first labeling, the primary antibodies used were: rabbit - IgG Anti-PV (Invitrogen, PA1-933), rabbit - IgG Anti-Sst (Invitrogen, PA5-87185), and rabbit - IgG Anti-Vip (Invitrogen, PA5-85616), with goat - IgG Anti-Rabbit - AlexaFluor-647 (Invitrogen, A32733) as the secondary antibody. In the second labeling, the primary and secondary antibodies used were: mouse - IgG2a Anti-GAD67 (Millipore, Mab5406), and goat - IgG2a Anti-Mouse -AlexaFluor-488 (Invitrogen, A21121).

### Morphological reconstructions

Biocytin-filled neurons were reconstructed manually using Neurolucida software (MBF Bioscience, Williston, VT, USA) attached to an Olympus BX50/51 or a Zeiss Axioplan microscope, equipped with a 100x oil immersion objective. The outlines of the pia surface, L4 barrels, and white matter were reconstructed on at low magnification (4x). These outlines were used to register each neuron to a standardized 3D reference frame of rat vS1 (43). After registration, we identified the somata of 9 INHs in L1, 53 INHs in L2/3, 99 INHs in L4, 74 INHs in L5 and 71 INHs in L6. To reconstruct the distributions of PV^+^, Sst^+^ and Vip^+^ INHs across vS1, a confocal laser scanning system (SP5, Leica Microsystems, Germany) with glycerol/oil immersion objectives (HC PL APO 10x 0.04 N.A., HC PL APO 20x 0.7 N.A.) was used to acquire first a single optical section at 10x from each of the 40-48 tissue sections (AlexaFluor-488: excitation at 488 nm, emission detection range: at 495-550 nm; AlexaFluor-647: excitation: 633 nm, emission detection range: 650-785 nm). The hence generated GAD67 images of each section were used to manually reconstruct the contours of the pia, L4 barrels, white matter, and blood vessels, the later were used to align the images from the 40-48 sections. We hence identified in each of the sections the location of the barrel column corresponding to the D2 whisker, and imaged each section again, now for a subvolume of 3 mm by 3 mm around the D2 column, for which we acquired high-resolution image stacks with a voxel size of 0.361 x 0.361 x 0.5 μm (20x glycerol objective with 2.0 digital zoom, 8x line average). The image stack for each 50 μm thick section hence comprised ∼100 optical sections. For each brain section, we aligned the 2D images (i.e., 1 optical section of the entire section at 10x) with the corresponding 3D images (i.e., ∼100 optical sections of 3×3 mm around the D2 column at 20x). In both image datasets, we manually identified all GAD67^+^ neurons in vS1, as well as all the GAD67^+^ INHs that co-expressed the respectively labeled molecular marker (PV, Sst, or Vip) using AmiraZIBEdition (65). The distributions obtained from the 2D images were scaled to match those obtained from the corresponding 3D images (66). The scaling factor β across *n* consecutive brain sections satisfied *y*_*i*_ = *βx*_*i*_ + *∈*, where {x_*i*_, y_*i*_}^*n*^ are the distributions in 2D and 3D images, respectively (for PV: β=3.8 (R^2^=0.98, p<10^−28^) and β=3.6 (R^2^=0.95, p<10^−18^), for Sst: β=3.6 (R^2^=0.94, p<10^−21^) and β=4.5 (R^2^=0.86, p<10^−15^), for Vip: β=5.6 (R^2^=0.85, p<10^−14^) and β=5.2 (R^2^=0.89, p<10^−15^). The distribution of molecular markers relative to their corresponding GAD67 distribution was robust across rats (two-sample KS between experiments: PV p=0.738, Sst p=0.545, Vip p=0.545). The distribution of -/-/- INHs was calculated as the difference between the GAD67 distributions (N=6) and the sum of the PV, Sst and Vip distributions (each N=2).

### Morphoelectrical classification

We recorded the firing responses to a series of 0.5-1 s long step current pulses, ranging from −100 pA to 500 pA. For analysis, we selected the voltage trace with an initial inter-spike-interval (ISI) closest to 100 Hz (36, 40, 67). These traces were standardized to 700 ms trial duration and resampled at 100 kHz (i.e., 100 ms before the step current, 500 ms step current, 100 ms after the step current). Thus, for recordings with current injections longer than 500 ms, APs after 500 ms were not considered for analysis. Traces with <3 APs were excluded. The electrophysiology was quantified similar to (2), including time-dependent E-features such as ISI shape and PSTH **(Table S1)**. The E-features were obtained from sparse principal component (SPC) analysis. 31 SPCs with explained variance (PEV) >1% were obtained. Two additional E-features were: (i) spiking frequency in Hz (i.e., number of APs divided by the time between first and last AP), and (ii) spiking adaptation in % (i.e., after AP frequencies were fitted to an exponential function, adaptation was defined as the difference between first and last fitted points normalized by the first fitted point). The morphology was quantified as suggested previously (2, 9, 22, 36–41), including barrel cortex-specific M-features such as the axon and dendrite inside and outside of the column. We extracted 47 M-features **(Table S2)**. The ME-features with low coefficients of variation (CV < 0.25) were removed. Only one from sets of highly correlated (r > 0.95) ME-features was selected. All ME-features were z-scored. An assignment module performed a preliminary clustering step for each modality, followed by a reduction of the number of clusters using an entropy criterion. Then, a sensing module evaluated cluster stability by resampling the data, followed by an entropy criterion to merge unstable clusters (66). The outputs are class assignments and performance metrics per modality **(Fig S2)**. We complement our dataset for rat vS1 with datasets reported for mouse V1 (10), mouse M1 (32), and human MTG (35). For mouse V1, we analyzed 528 INHs with reconstructed morphologies, of which 466 had electrophysiological recordings with at least 3 APs, and 511 had assigned transcriptomic cluster IDs. The molecular type of a given INH was determined from their corresponding T-cluster. Thus, 466 INHs had data available across all three modalities. For mouse M1, we analyzed 370 INHs with reconstructed morphologies, of which 363 had electrophysiological recordings, and 368 had assigned T-cluster IDs. In human MTG, we analyzed 140 INHs from acute brain and cultured slice preparations. We excluded recordings from cultured slices, because electrophysiological, but not morphological, properties differ significantly between the two conditions (35). Accordingly, among the 140 INHs with reconstructed morphologies, 59 had electrophysiological recordings from acute slices, and 136 had assigned T-cluster IDs. We also analyzed 173 EM-reconstructed INH morphologies from the MICrONS dataset (34) – i.e., from all cortical layers of visual cortex tissue (primary and higher-order areas) from a male mouse aged 87 days. INHs were assigned to morphological types (68) as follows: basket cells targeted >20% of their synapses to somata or proximal dendrites, Martinotti cells targeted distal dendrites of pyramidal neurons without features of bipolar or neurogliaform cells, bipolar cells had two to three primary dendrites and mainly targeted other INHs, and neurogliaform cells had sparse outputs and boutons with vesicles but no clear postsynaptic targets. CAVEclient (69) with datastack minnie65 public was used for materialization version 1621 (Nov 2025). The `aibs_metamodel_celltypes_v661` table was accessed to obtain automated M-types. Ref. (51) utilized `pt_root_id` in materialization version 765 to identify the INHs analyzed in this study. As `pt_root_id` may change across versions, we used a snapshot from September 9, 2023 at 4:00 AM to query the `nucleus_detection_v0` table and determine their corresponding `nucleus_id`, which remains unchanged across materialization versions. We hence assigned M-types to the 173 EM-reconstructed INHs.

### Analysis of clustering results

We generated the ME-diversity space by unfolding the UMAP along the soma depth dimension by affine transformations **(Fig. S4B)**. The UMAP algorithm used a local neighborhood of size 50, a minimum distance between points of 0.2, and a cosine similarity metric. We unfolded the UMAP as follows: (i) each INH in the UMAP was mapped to a closed unitary ball around the origin given by *B* = {*x* ∈ ℝ^*n*^: ‖*x*‖_2_ ≤ 1}, (ii) the INHs in this normalized UMAP were clustered into L1-3, L4 and L5-6 INHs, using a Gaussian mixture model with three components, (iii) L1-3 and L4 INHs were flipped vertically and horizontally in the UMAP, respectively, to align them with the L5-6 INHs, (iv) a linear regression was applied to obtain the angles with respect to the origin for L1-3, L4 and L5-6 INHs, respectively, (v) the angles and an inverse rotation matrix were used to place L1-3, L4 and L5-6 INHs along the horizontal axis, (vi) L1-3, L4 and L5-6 INHs were centered at the origin, and (vii) L1-3 and L5-6 INHs were shifted symmetrically along the horizontal axis. Gradients were calculated by fitting a linear regression between the spatial coordinates of the ME-diversity space and the z-scored values of the ME-feature. Groups I-IIIa/b were obtained by clustering INHs binned by soma depth (i.e., depth-independent grouping). The bin size was set to 900 μm, resulting in two groups, which covered approximately L1-4 (160 INHs) and L5-6 (139 INHs). We performed our clustering procedure with a hierarchical algorithm (70) and assigned the resulting 67 clusters to Groups I-III: at any cortical depth, INHs with highest spiking frequencies (z-score>0.15) and weakest adaptation (z-score<−0.35) were Group I; INHs with lowest spiking frequencies and strongest adaptation were Group II (spiking frequency < 3.0 · adaptation + 0.05); INHs with small vertical axon extents and large vertical dendrite extents were Group IIIa (axon < 0.15 · dendrite + 0.35); INHs with low spiking frequencies (z-score<−1) or smallest vertical dendrite extents (z-score<−0.5) were Group IIIb. The distributions of Groups I-IIIa/b **(Fig. 5E)** represent the respective numbers of INHs in our sample at 50 μm cortical depth resolution, normalized the total number of INHs at each depth. We smoothed these depth-profiles independently for each group with a moving average of 200 μm in size.

### Quantification and statistical analysis

All data are reported as mean ± 95% confidence interval (CI) unless stated otherwise. All statistical details can be found in the main text and figure legends, including the statistical tests used, sample size, and what the sample size represents (e.g. number of animals, number of cells). Significance was defined for p-values smaller than 0.05. Boxplots are median, 25^th^/75^th^ percentiles and whiskers extend up to 1.5 times the interquartile range. All data beyond the whiskers are shown as outliers.

## Acknowledgments

We thank Staci Sorensen for providing the data from mouse V1 and Daniel Udvary for a first analysis of the vS1 data. Funding was provided by the Max Planck Institute for Neurobiology of Behavior – caesar (MO), the Max Planck Institute for Biological Cybernetics (MO), the Max Planck Florida Institute for Neuroscience (HM, MO, BS), the Helmholtz Society (DF), European Research Council grant 633428 (MO), Deutsche Forschungsgemeinschaft grants SFB 1089 (MO), SPP 2041 (MO) and Fe472/2-1 (DF), German Federal Ministry of Education and Research grants 01GQ1002 and 01IS18052 (MO), Neuroscience Network North Rhine-Westphalia grant iBehave (MO), NIH grant R24MH117295 (ND), and a NeuroData Discovery Award from the Kavli Foundation (FY).

## Author Contributions

FY, BS and MO conceived and designed the study. FY analyzed data, discovered the shifts of morphoelectric properties with cortical depth, and wrote the first draft of the paper. GQ, HM, DF, BS and MO provided the vS1 data. FM performed immunolabeling experiments. ND contributed to the analysis of V1, M1 and MTG data. MO wrote the final draft of the paper with FY and DF.

## Supplementary Figures

**Figure S1.**
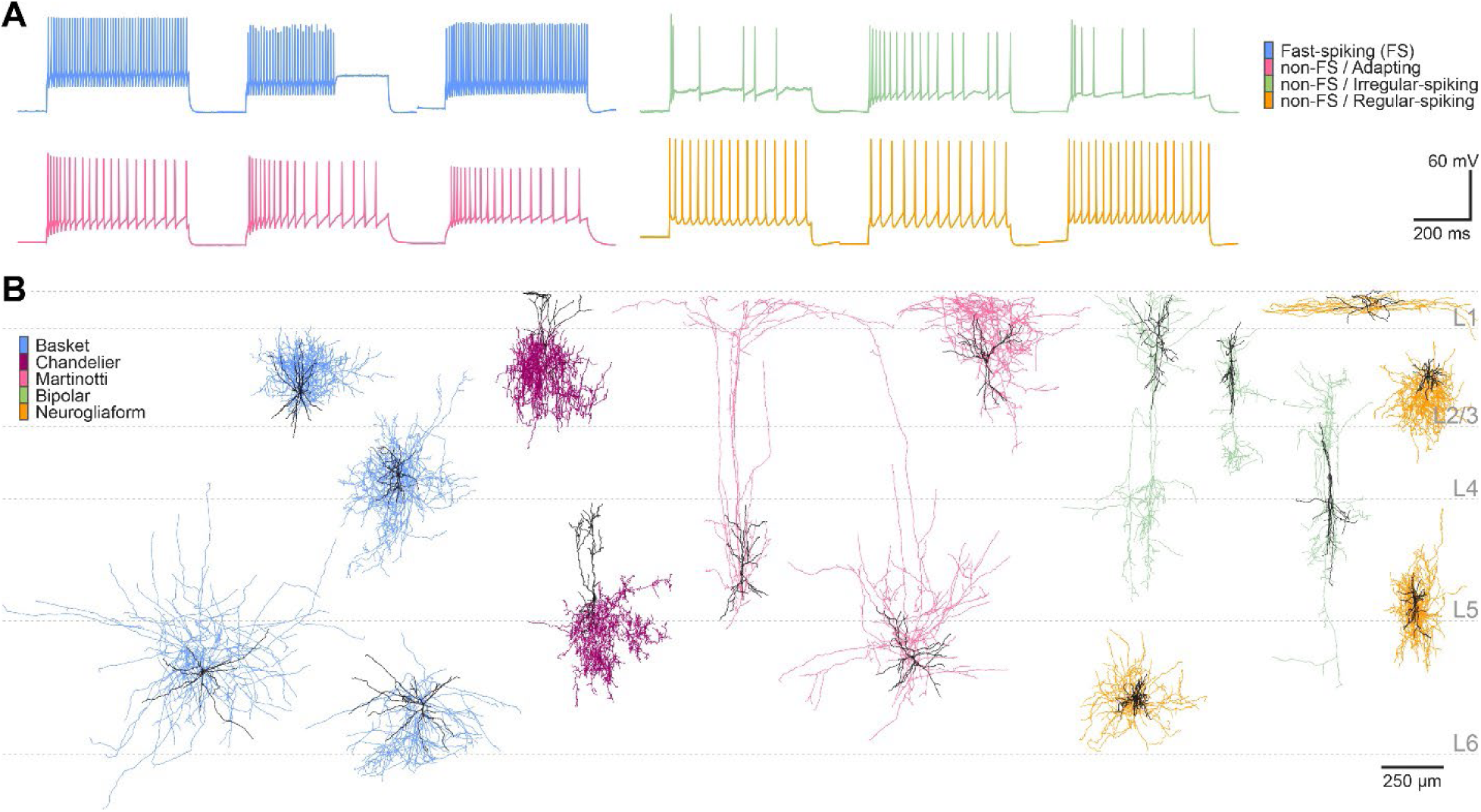
Electrophysiological and morphological INH types in our sample. **(A)** Voltage trace examples for the major E-types of cortical INHs, i.e., fast-spiking (FS) versus non-FS INHs, and the latter with adapting, irregular or regular spiking patterns. **(B)** Example morphologies for major M-types of cortical INHs, i.e., basket, chandelier, Martinotti, bipolar, and neurogliaform cells. Axons are colored by M-type, and dendrites are colored in black.

**Figure S2.**
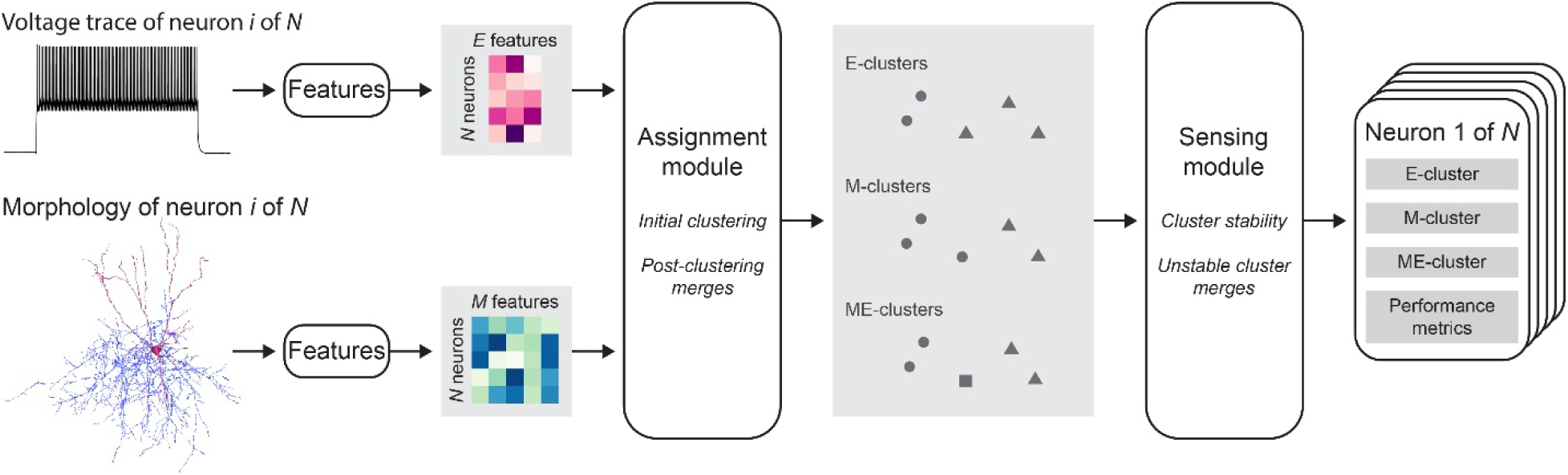
Unsupervised multimodal clustering method. Features are calculated from a given registered INH morphology and voltage trace. The assignment module performs an initial clustering step dependent on the modality, followed by a reduction of the number of clusters using an entropy criterion. The sensing module evaluates cluster stability, followed by an entropy criterion to merge unstable clusters. The output is the final cluster assignment and performance metrics per modality.

**Figure S3.**
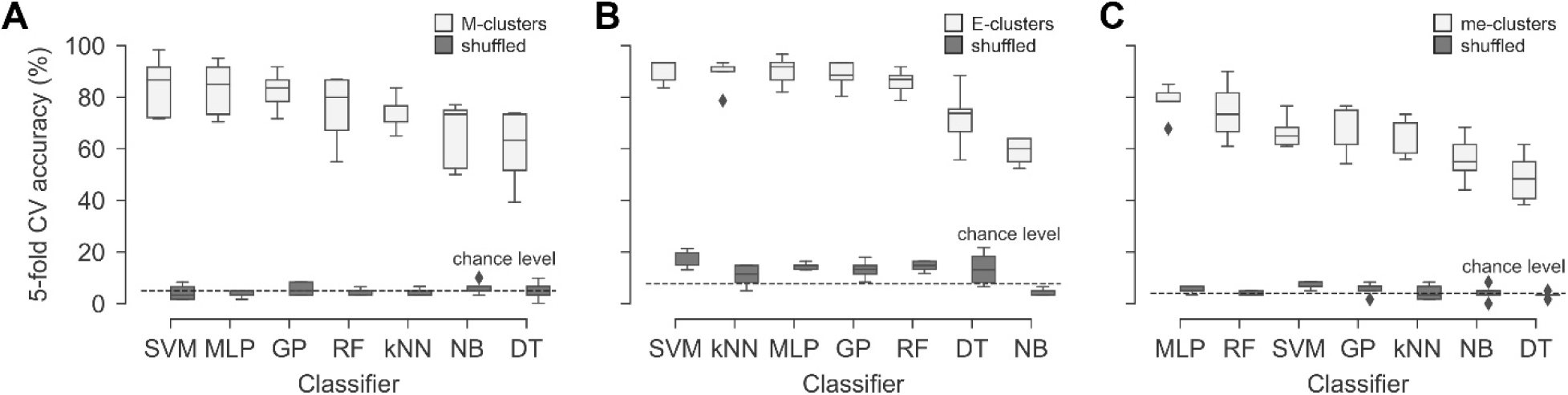
Validation of final cluster assignments via 5-fold cross-validation (CV). **(A)** Seven different classifiers were utilized: a linear support vector classifier (SVM), a multi-layer perceptron classifier with α = 1 and 1000 maximum iterations (MLP), a Gaussian process classifier (GP), a random forest classifier with 500 estimators (RF), a k-nearest neighbors classifier with k = 5 (kNN), a naive Bayes classifier (NB), and a decision tree classifier with a maximum tree depth of 500 (DT). Chance level is the inverse of the number of M-clusters, i.e., 5.0%. The results utilizing the original cluster assignments (m-clusters) as labels are shown in a lighter shade of gray, whereas with shuffled assignments in a darker shade. We calculated the out-of-bag prediction accuracy of a RF classifier (79.5%), and the leave-one-out CV of a SV classifier (88.1%) and compared them against the results from Gouwens et al. (70% and 79%, respectively). **(B)** Same as panel A, but with two differences in the parameter settings: 1) MLP had a regularization parameter of α = 1.1, and 2) kNN used k = 3. Chance level is set to 7.7%. **(C)** Same as panel A, but with one difference in the parameter settings: kNN used k = 8. Chance level is set to 4.0%.

**Figure S4.**
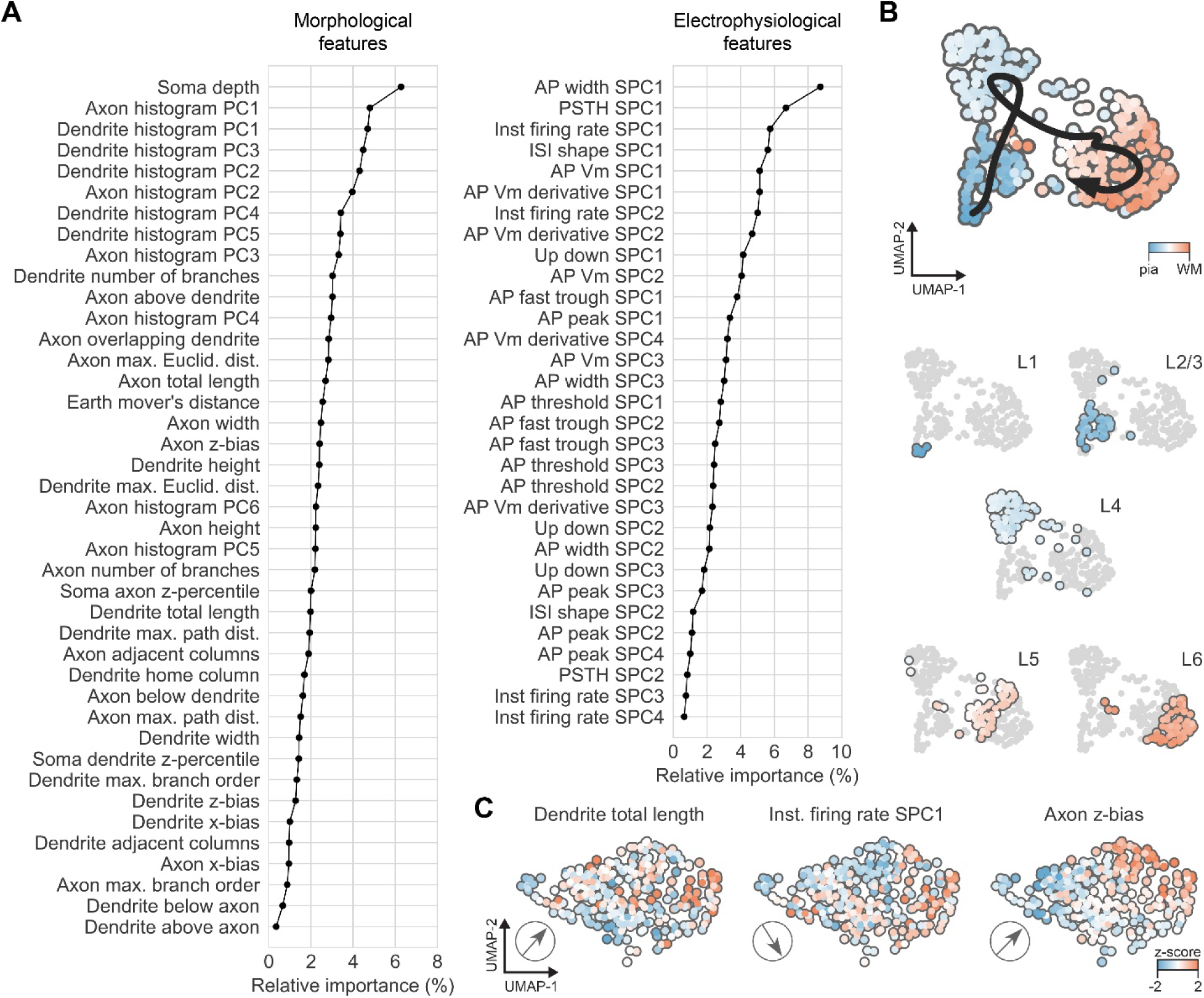
Importance of ME-features for INH clusters. **(A)** Gini impurity index of morphological (left) and electrophysiological (right) features used to determine M- and E-clusters, respectively. **(B)** Top: We unfolded the UMAP of the ME-feature space along the cortical depth dimension, which we approximated by fitting a spline to the UMAP at a resolution of 100 μm (see **Methods** for details). Bottom: INHs colored by their soma depth in the original UMAP shows that INHs from the same cortical layer are located close to one another in the feature space that give rise to the 25 ME-clusters. **(C)** Additional examples of features that display diagonal gradients in the ME-diversity space. The black arrows represent the gradients of these features across the ME-diversity space – i.e., pia is left and WM is right.

## Supplementary Tables

**Table S1.** Electrophysiological features. Dimensionality reduction was applied independently to each dataset, obtaining 2-4 sparse principal components (SPCs) with an adjusted explained variance larger than 1%. When applicable, voltage and time units are in mV and ms, respectively.

| Dataset name | # features | Definition |
| --- | --- | --- |
| AP $V_m$ | 3 | Membrane potential ($V_m$) of first action potential (AP). The time window is 3 ms (with a sampling rate of 100 kHz): 1 ms before the peak of the AP and 2 ms after it. |
| AP $dV_m/dt$ | 4 | Time derivative of AP $V_m$. |
| ISI shape | 2 | Average of inter-spike-interval (ISI) voltage trajectories, linearly interpolated in time to fit a standardized query of 100 points. |
| PSTH | 2 | Peri-stimulus time histogram (PSTH) normalized by bin width, with a bin width of 50 ms. |
| Inst. firing rate | 4 | Instantaneous firing rate as the inverse of each ISI. |
| AP peak* | 4 | For each AP, the maximum $V_m$ value. |
| AP threshold* | 3 | For each AP, the $V_m$ value where $dV_m/dt$ between pre-AP voltage and AP upstroke is maximized. The pre-AP voltage is the $V_m$ value at half-ISI before AP (for first AP, is the half-interval between stimulus onset and AP). The AP upstroke is the maximum value of $dV_m/dt$ between pre-AP voltage and AP peak. |
| AP fast trough* | 3 | For each AP, the minimum $V_m$ value between AP peak and post-AP voltage, where post-AP voltage is the $V_m$ value at half-ISI after AP (for last AP, is the half-interval between AP and stimulus offset). |
| AP width | 3 | For each AP, the width at half-height. |
| U/D-stroke ratio | 3 | For each AP, the ratio of AP upstroke to AP downstroke, where AP downstroke is the minimum value of $dV_m/dt$ between AP peak and AP fast through. |

\* indicates that calculated values were placed at spike times and binned using a regular time window of 20 ms. Values falling in the same bin were averaged. Zero valued bins between non-zero valued bins were linearly interpolated.

**Table S2.** Morphological features. The pia to white matter direction is denoted as *z*, whereas the dorsal medial to ventral lateral direction as *x*. When applicable, length units are in mm.

| Category | Feature name | # features | Definition |
| --- | --- | --- | --- |
| Location | Soma depth | 1 | Distance between the registered pia and soma. |
| Length | Total length* | 2 | Length across all branches. |
|  | {Height, width}* | 4 | Extent in the {z,x}-direction. |
| Bias | {z,x}-bias* | 4 | Difference in extent in {z,x} in one direction from the soma and the other. Values are signed for z-bias and unsigned for x-bias. |
| Branch | Max. branch order* | 2 | Maximum number of bifurcations encountered between the soma and all neurite tips. |
|  | Number of branches* | 2 | Number of individual branches in the morphology. |
| Distance | Max. Euclidean distance* | 2 | Direct-line distance from the soma to the furthest neurite tip. |
|  | Max. path distance* | 2 | Path distance from the soma to the furthest neurite tip. |
|  | Max. mean contraction* | 2 | Average of the ratios of the summed Euclidean distance between bifurcations, and between bifurcations and tips, to the summed path distance between same. |
| Overlap | Axon or dendrite {above, overlapping, below} dendrite or axon | 6 | Fraction of axon or dendrite nodes {above, overlapping, below} the full z-extent of the dendrite or axon, respectively. |
|  | EMD | 1 | Earth mover's distance metric calculated between the normalized and aligned axon and dendrite depth profile histograms. |
| Percentile | Soma {z,x}-percentile* | 4 | Percentile location of the {z,x}-coordinate of the soma within the {z,x}-coordinates distribution of all nodes. |
| Depth histogram | Depth histogram PCs* (6 axon and 5 dend.) | 11 | Principal components (PCs) produced by PCA to depth profile histograms aligned to our reference frame (50 um bins) that exceed 5% explained variance. |
| Column | Home column* | 2 | Relative length within the D2 column. |
|  | Adjacent columns/septa* | 2 | Relative length outside the D2 column. |

\* indicates that the feature was calculated independently for the axon and dendrite morphology.

**Table S3.** Molecular composition of the rat barrel cortex (vS1).

| Whisker | Volume | GAD67 | PV | Sst | Vip |
| --- | --- | --- | --- | --- | --- |
| α | 0.14 ± 0.02 | 1479 ± 6 | 631 ± 539 | 251 ± 136 | 433 ± 220 |
| β | 0.17 ± 0.02 | 1917 ± 759 | 675 ± 404 | 296 ± 103 | 557 ± 73 |
| γ | 0.25 ± 0.08 | 1867 ± 913 | 1202 ± 746 | 355 ± 286 | 797 ± 33 |
| δ | 0.20 ± 0.04 | 2577 ± 1096 | 799 ± 295 | 352 ± 49 | 749 ± 251 |
| A1 | 0.14 ± 0.04 | 2140 ± 865 | 744 ± 564 | 203 ± 125 | 425 ± 63 |
| A2 | 0.14 ± 0.02 | 2124 ± 952 | 686 ± 208 | 259 ± 73 | 323 ± 58 |
| A3 | 0.10 ± 0.01 | 1535 ± 1190 | 442 ± 278 | 185 ± 49 | 241 ± 178 |
| A4 | 0.10 ± 0.02 | 1760 ± 1048 | 516 ± 584 | 144 ± 120 | 289 ± 189 |
| A-row | 0.12 ± 0.02 | 1890 ± 430 | 597 ± 190 | 198 ± 48 | 319 ± 72 |
| B1 | 0.16 ± 0.01 | 1933 ± 896 | 608 ± 286 | 286 ± 259 | 382 ± 127 |
| B2 | 0.17 ± 0.03 | 2075 ± 1068 | 764 ± 481 | 339 ± 252 | 368 ± 82 |
| B3 | 0.15 ± 0.03 | 2068 ± 669 | 568 ± 183 | 317 ± 296 | 252 ± 33 |
| B4 | 0.15 ± 0.02 | 1979 ± 601 | 635 ± 538 | 276 ± 31 | 266 ± 129 |
| B-row | 0.16 ± 0.01 | 2014 ± 317 | 644 ± 160 | 304 ± 90 | 317 ± 57 |
| C1 | 0.21 ± 0.03 | 1935 ± 50 | 635 ± 203 | 490 ± 383 | 500 ± 142 |
| C2 | 0.21 ± 0.03 | 2117 ± 561 | 802 ± 294 | 489 ± 393 | 540 ± 215 |
| C3 | 0.23 ± 0.02 | 2221 ± 662 | 951 ± 387 | 464 ± 143 | 491 ± 132 |
| C4 | 0.23 ± 0.04 | 2398 ± 183 | 1033 ± 699 | 550 ± 201 | 502 ± 22 |
| C-row | 0.22 ± 0.02 | 2162 ± 227 | 855 ± 200 | 498 ± 116 | 508 ± 57 |
| D1 | 0.21 ± 0.03 | 1786 ± 520 | 937 ± 687 | 447 ± 314 | 432 ± 19 |
| D2 | 0.28 ± 0.04 | 2464 ± 1194 | 1420 ± 669 | 616 ± 304 | 643 ± 59 |
| D3 | 0.26 ± 0.04 | 2163 ± 697 | 1162 ± 304 | 636 ± 201 | 747 ± 411 |
| D4 | 0.24 ± 0.03 | 3060 ± 451 | 1212 ± 75 | 480 ± 201 | 460 ± 100 |
| D-row | 0.25 ± 0.02 | 2368 ± 452 | 1183 ± 229 | 545 ± 116 | 571 ± 126 |
| E1 | 0.32 ± 0.04 | 3320 ± 801 | 1243 ± 520 | 672 ± 170 | 890 ± 143 |
| E2 | 0.35 ± 0.07 | 4337 ± 513 | 1408 ± 148 | 866 ± 241 | 954 ± 792 |
| E3 | 0.36 ± 0.05 | 3672 ± 751 | 1821 ± 274 | 685 ± 516 | 1092 ± 395 |
| E4 | 0.28 ± 0.04 | 3142 ± 175 | 1366 ± 403 | 652 ± 451 | 720 ± 258 |
| E-row | 0.33 ± 0.03 | 3618 ± 410 | 1460 ± 212 | 719 ± 155 | 914 ± 202 |
| Mean | 0.21 ± 0.01 | 2332 ± 222 | 928 ± 119 | 430 ± 64 | 544 ± 74 |

Displayed numbers are mean ± 95% CI (n = 2). Volume units are given in mm<sup>3</sup>.

